# A functional cortical network for sensorimotor sequence generation

**DOI:** 10.1101/783050

**Authors:** Duo Xu, Yuxi Chen, Angel M. Delgado, Natasha C. Hughes, Mingyuan Dong, Linghua Zhang, Daniel H. O’Connor

## Abstract

The brain generates complex sequences of movements that can be flexibly reconfigured in real-time based on sensory feedback, but how this occurs is not fully understood. We developed a novel ‘sequence licking’ task in which mice directed their tongue to a target that moved through a series of locations. Mice could rapidly reconfigure the sequence online based on tactile feedback. Closed-loop optogenetics and electrophysiology revealed that tongue/jaw regions of somatosensory (S1TJ) and motor (M1TJ) cortex encoded and controlled tongue kinematics at the level of individual licks. Tongue premotor (anterolateral motor, ALM) cortex encoded intended tongue angle in a smooth manner that spanned individual licks and even whole sequences, and progress toward the reward that marked successful sequence execution. ALM activity regulated sequence initiation, but multiple cortical areas collectively controlled termination of licking. Our results define a functional cortical network for hierarchical control of sensory- and reward-guided orofacial sequence generation.

---

The world presents itself to us as a series of sensations that typically arises from our own actions, and in turn elicits further actions, forming an intricate sensorimotor loop. A desired sequence frequently cannot be achieved by chaining together a series of “reflexes” (or simple sensory-motor pairs) ^1^ because sensory inputs may not capture a changing environment at a particular moment in time, but the motor output is still required to change. The brain therefore must keep a memory of past states and guide ongoing actions in part by inference. Memorized motor sequences, however, must be flexibly rearranged or branched upon arrival of relevant sensory input. Sensorimotor sequences may also exhibit a hierarchical organization where elementary patterns of sensory input and motor kinematics form short sequences that are structured into longer and more complex compositions ^2^. As many sequences are learned, it is also critical to understand interactions between motor control and reward signals ^3^. A fascinating and unresolved question, therefore, is how the brain organizes such hierarchical representation and flexible control.

Sensorimotor control of orofacial structures governs a wide range of both innate and learned behaviors that are essential for survival, such as communication, breathing and feeding, and is thus tightly orchestrated ^4, 5^. The motor control of directional licking has been studied extensively in mice and yielded many insights into working memory, motor planning, and the neural basis of simple sensorimotor transformations ^6–13^.

Here, we show that mice can learn to perform flexible feedback- and memory-guided movements composed of individual licks that are organized into distinct sequences. Mice used their tongue to sequentially reach a set of predefined targets with remarkable speed and accuracy. When the target “backtracked” to a prior position unexpectedly, mice used tactile feedback to modify the ongoing sequence and quickly relocate the target. Mice alternated between two sequences across trials in the absence of external cues. This type of flexible feedback control, although common in daily life, differs from typical cerebellum based sensorimotor adaptation, where movements are fine tuned based on sensory prediction errors ^14, 15^. The necessity of sensory feedback for ongoing execution also distinguishes our task from those involving repetitive or non-dexterous movement sequences ^16, 17^.

Closed-loop optogenetic inhibition and population single-unit electrophysiological recordings from multiple brain areas allowed us to identify three main cortical regions involved in controlling the sensorimotor sequences. The tongue premotor region (anterior lateral motor cortex, ALM) ^9, 18^ was critical in initiating a sequence and controlling intended tongue angle smoothly across a series of individual licks. ALM neurons encoded sequence direction, progress toward the reward that signaled sequence completion, and intended tongue angle for upcoming sequences during the inter-trial interval, supporting a memory for how sequences were ordered within the behavioral session. In contrast, activity in the tongue/jaw region of primary motor cortex (M1TJ) strongly encoded instantaneous lick angle, tongue length and tongue velocity. Tongue/jaw primary somatosensory cortex (S1TJ) activity encoded tongue angle, length and velocity, and its inhibition randomized lick angles, suggesting a proprioceptive role for these signals in S1TJ. Termination of licking depended on activity in multiple cortical regions.

Our results reveal a functional sensorimotor cortical network that allows mice to use the tongue to perform complex, flexible and sensory-guided motor sequences.

## Results

### Sequence licking task

We trained head-fixed mice to perform a task in which they used sequences of directed licks to advance a motorized port through 7 consecutive positions, either from left to right or right to left, after an auditory cue (15 kHz, 0.1 s) that signaled the start of a trial (Fig. 1a; Movie 1). Each transition from one position to the next was driven in a closed-loop manner by a single lick touching the port. Thus, if a lick missed the port, the port would remain at the same position until the tongue eventually made contact. The port was no longer movable after the mouse had finished the 7 positions and a water droplet was delivered as a reward after a short delay (0.25 s, or 0.5 s in two mice). The next trial would then start after a random inter-trial interval (ITI) with a mean duration of 6 s, and the sequence would go in the opposite direction.

**Fig. 1.**
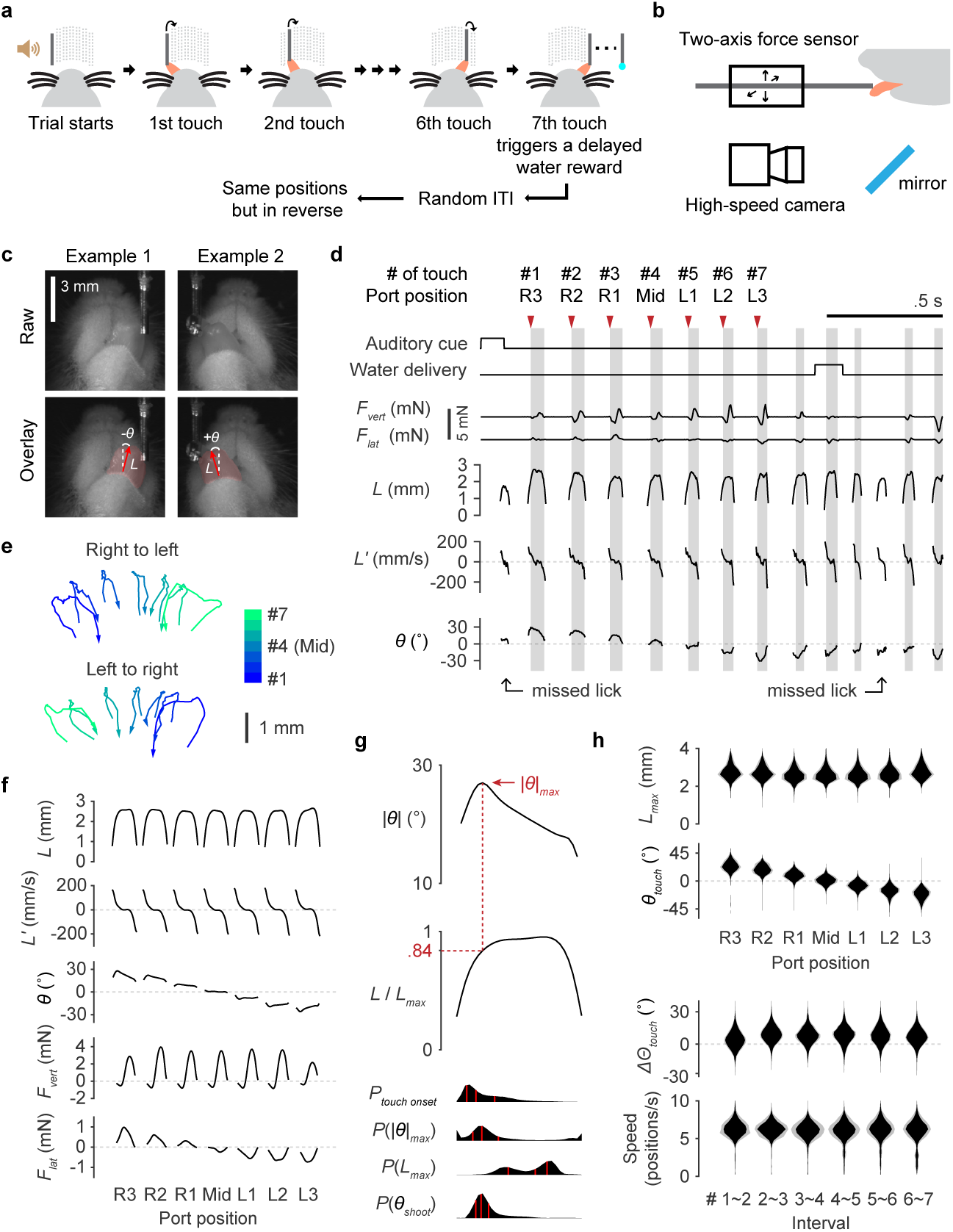
Sequence licking task. **a**, Schematic of the sequence licking task. **b**, Schematic of the two-axis optical force sensors and high-speed videography in relation to a head-fixed mouse. **c**, Zoomed in high-speed video images showing the bottom view of a mouse licking the port. Overlaid vectors in red are outputs from the regression DNN, which point from the base to the tip of the tongue. Tongue length (*L*) is defined by the vector length. Tongue angle (*θ*) is the rotation of the vector from midline (positive being to the mouse’s right). Light red shading depicts shape of the tongue based on output from the segmentation DNN. **d**, Time series of task events and behavioral variables during the first 2 s of an example trial. Variables recorded from the force sensors include the vertical lick force (*F_vert_*, positive acts to lift the port up) and the lateral lick force (*F_lat_*, positive acts to push the port to the right). Kinematic variables including *L*, its rate of change (*L’*) and *θ* were derived from high-speed video. Periods of tongue-port contact are shaded in gray and are numbered (#) sequentially. R3, R2, R1, Mid, L1, L2 and L3 indicate the 7 port positions from the rightmost to the leftmost. Arrowheads indicate the touch onsets for which a port movement (or the delayed water delivery at the last position) was triggered. **e**, Two example trials showing the trajectories of the tongue tip when a mouse sequentially reached the 7 port positions, for both sequence directions. Arrows indicate the direction of time within each trajectory. **f**, Patterns of kinematics and forces of single licks at each port position (n = 22003 trials; mean ± 95% bootstrap confidence interval). The duration of individual licks was normalized. **g**, Top, the pattern of angle deviation from midline (|*θ* - 0°| or simply |*θ*|) of single licks at R3 and L3. The vertical line indicates maximum |*θ*| (|*θ*|*_max_*). Middle, tongue length (*L*) expressed as a fraction of its maximum (*L_max_*). The horizontal line indicates, on average, the fraction where |*θ*|*_max_* occured. Bottom, time aligned probability distributions showing when touch onset, |*θ*|*_max_*, *L_max_* or *θ_shoot_* occured. Red lines mark quartiles. n = 22003 trials. Lick patterns show mean ± 95% bootstrap confidence interval. **h**, Top, probability distributions of *L_max_* and *θ_touch_* for licks at each port position. Bottom, probability distributions of the change in *Θ_touch_* (*ΔΘ_touch_*) and instantaneous sequence speed (Methods) for each interval separating port positions. Distributions show mean ± SD across n = 15 mice.

We used high-speed (400 Hz) video to capture tongue motions during sequence performance (Fig. 1b) and developed deep artificial neural networks (DNN) to extract horizontal tongue angle (*θ*) and length (*L*) from each video frame (Methods; Fig. 1c, Extended Data Fig. 1a,b). To quantify the sensory feedback experienced by the tongue when touching the port, we developed a set of sensors that measured the vertical and lateral component of instantaneous contact force (*F_vert_* and *F_lat_*) (Fig. 1b, Extended Data Fig. 1c,d). Electrical contact detection was used to determine the precise onset and offset times of touch. An example trial shows all the measurements aligned in time (Fig. 1d).

Mice modulated each lick differently to reach different target locations (Fig. 1e). Specifically, the modulation was mainly in *θ* whereas the patterns of *L* and its rate of change (*L’*) remained similar across targets (Fig. 1f). When focusing on a single pattern at most lateral positions, we saw the tongue shooting out and quickly, but only briefly, reaching maximal deviation from midline (*|θ* - 0°*|_max_* or simply *|θ|_max_*) (Fig. 1g). As a result, the onset of touch mostly occurred around *|θ|_max_*. Interestingly, the modulation of *L* did not match that of *|θ|*, suggesting a potential dissociation of control. Later, when analyzing licks which may or may not have contact, we use *θ_shoot_*, defined as the *θ* when *L* reaches 0.84 maximal *L* (*L_max_*), to succinctly depict the lick angle (Fig. 1g). In addition, we will use capital *Θ* to represent unified tongue angles where the sign in right to left sequences is flipped so that data with both sequence directions can be pooled together.

Mice performed the task in darkness such that there were no visual cues to guide the licks. We played both white noise and pre-recorded mechanical noise of port transitions as masking sounds to prevent mice from using auditory cues (Methods). In addition, temporarily induced hearing loss (Methods) rendered mice unable to respond to the auditory cues but did not affect sequence performance, except in one of five mice, compared with control (Extended Data Fig. 1e). To test whether or not mice used odor emanating from the port for localization, we masked out any potential odor by applying a constant flow (2 LPM) of fresh air covering the nose and port region (Methods), and found no significant drop in sequence performance, except in one of six mice, compared with the condition without air flow (Extended Data Fig. 1f).

Mice typically obtained proficiency in standard sequences after a total of ∼1500 trials of training (Methods; Extended Data Fig. 1g,h). In addition to stereotypic licking kinematics, expert mice showed remarkable speed of sequence execution, with the 7 positions completed in about a second (Fig. 1h).

### Flexible execution of sequences with backtracking

To determine if the sequence generation after training was strictly “ballistic” or was capable of flexible reconfiguration based on sensory feedback, we next varied the task by introducing unexpected port transitions after mice had learned the standard sequences (Fig. 2a; Movie 2). Specifically, in a randomly interleaved subset (1/3 or 1/4) of trials, when a mouse licked at the middle position, the port would backtrack two steps rather than continue to the anticipated position. Mice previously trained only with standard sequences learned (Methods; Fig. 2b, Extended Data Fig.1i,j) to detect the change of port transition, lick to the new position and finish the rest of the sequence. On average, it took 1 to 2 missed licks before mice quickly relocated the port (Fig. 2c).

**Fig. 2.**
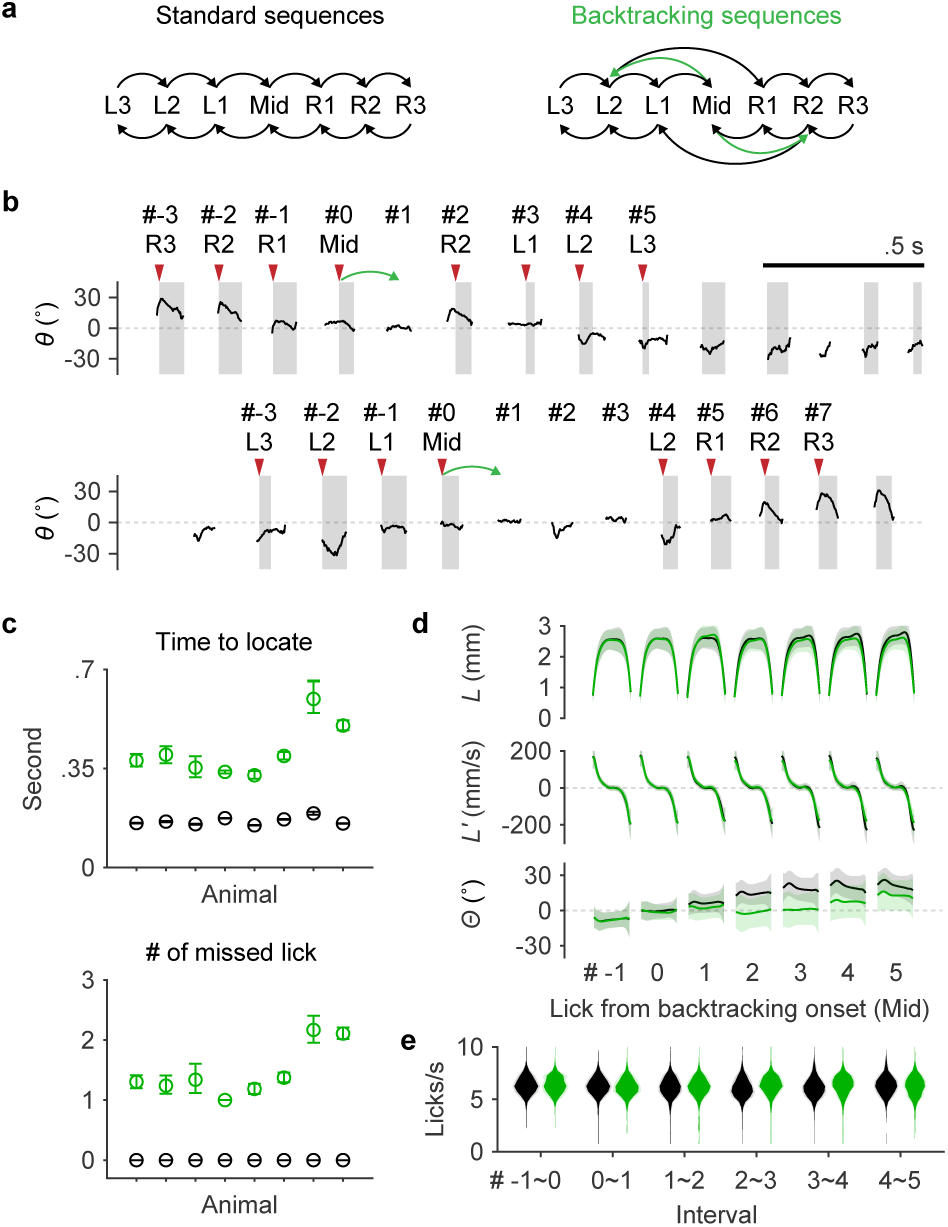
Flexible execution of sequences with backtracking. **a**, Transition diagrams depicting standard sequences and those with backtracking (green). **b**, Example trials where the port backtracked (green arrows) when a mouse touched Mid. Licks including both touches and misses are indexed with respect to the lick at Mid. Top, the mouse missed once before it successfully relocated the port and finished the rest of the sequence. Bottom, after an initial miss and a lick back, the mouse made a second attempt and caught the port. **c**, Top, time to locate the port at its next position during the 4th interval, for standard sequences (black) or for sequences when the port backtracked (green). Bottom, the number of missed licks during the 4th interval. Mean ± 95% bootstrap confidence interval. **d**, *L*, *L’* and *Θ* patterns for seven consecutive licks aligned at the 4th (Mid) touch (#0). Licks in standard sequences (n = 7365 trials) are shown in black, those in backtracking sequences (n = 2674 trials) are in green. Mean ± SD. **e**, Probability distributions of instantaneous lick rate for each interval separating consecutive pairs of the seven licks during standard (black) or backtracking (green) sequences (n = 8 mice; mean ± SD).

The lick patterns of *L* and *L’* appeared identical whether a mouse was performing a standard sequence or one with backtracking (Fig. 2d). In contrast, *Θ* was strongly modulated toward backtracked directions starting at the second lick after backtracking occurred. A slight change in *Θ* was also present in the first lick after backtracking, presumably due to the mouse sensing the onset of port motion on its tongue toward the end of the prior lick. To our surprise, mice did not pause the ongoing sequence after detecting backtracking (Fig. 2e). Instead, they showed the same, if not higher, lick rate when making adjustments in single lick bouts. This suggests that flexible control of sequence execution can be independent from the pattern generators responsible for generating basic rhythmic licking patterns ^19^.

Together, these observations demonstrate that head-fixed mice can learn to perform complex and flexible licking sequences guided by sensory feedback.

### Closed-loop optogenetic inhibition screen of cortical areas

To determine which brain regions contributed to the performance of our novel sequence licking task, and at which points during sequence execution, we performed systematic closed-loop optogenetic silencing experiments.

We used the “clear-skull” preparation ^9^, a method that greatly improves the optical transparency of intact skull, to non-invasively photoactivate channelrhodopsin-expressing GABA-ergic neurons and thus indirectly inhibit nearby excitatory neurons (Fig. 3a). In different experimental sessions, bilateral inhibition was centered at each of five regions: the ALM ^20^ cortex, a somatomotor region centered at the body-related primary motor cortex (M1B) ^21, 22^, the S1TJ cortex ^13, 23^, the macrovibrissae subregion (or barrel field) of the primary somatosensory cortex (S1BF), and the trunk subregion of the primary somatosensory cortex (S1Tr, also including a part of posterior parietal cortex). For each region, inhibition was triggered with equal probability (10%) at sequence initiation, at mid-sequence or at the start of water consumption (Fig. 3b). Importantly, stimulation at mid-sequence and at consumption was triggered in closed-loop by the onset of the fourth (middle) touch during the sequence and by the onset of the first touch after water delivery, respectively. In the case of consumption, the trigger conditions ensured that the tongue had contacted water before inhibition started.

**Fig. 3.**
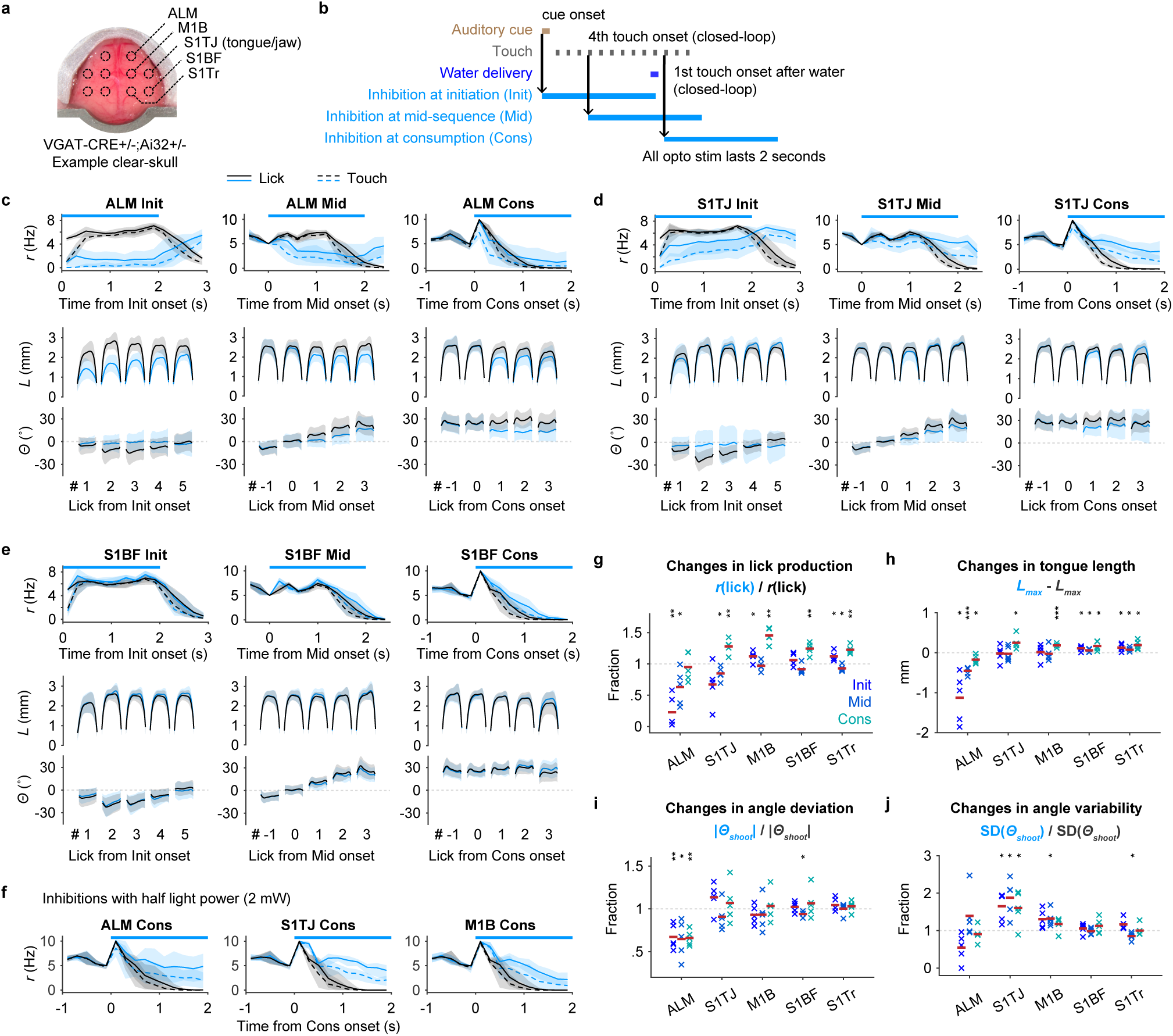
Closed-loop optogenetic inhibition defines cortical areas involved in sequence control. **a**, Example clear-skull preparation overlaid with circles indicating the five bilateral target regions. **b**, Triggering scheme for photoinhibition at sequence initiation, mid-sequence and water consumption. **c**, Effects of ALM inhibition for each of the three periods. Top row, the rate of licks (solid lines) or touches (dashed lines) as a function of time for trials with (blue) or without (black) inhibition (mean ± 95% hierarchical bootstrap confidence interval). Horizontal bars indicate the 2 s long illumination. Time zero is the onset of inhibition, or the equivalent time in trials without inhibition. Middle and bottom rows, *L* and *Θ* patterns (mean ± SD) for five consecutive licks in trials with (blue) or without (black) inhibition. Lick #0 is the lick that triggered inhibition, #1 the first lick after the trigger, #-1 the lick before, etc. Licks from inhibited trials are colored blue and those without inhibition are in black. n = 1181 trials from 5 sessions, one session per mouse. **d-e**, Same as (**c**) but for S1TJ (n = 1542 trials) and S1BF (n = 1399 trials). **f**, Effects of low power (2 mW) inhibition at ALM (n = 850 trials), S1TJ (n = 774) and M1B (n = 784) during consumption period. Same conventions as (**c**, right). **g**, Changes in lick production across regions and periods, quantified as lick rate in trials with inhibition as a fraction of lick rate in trials without inhibition. Red bars indicate means (n = 5 mice). ∗ p < 0.05, ∗∗ p < 0.01, ∗∗∗ p < 0.001, not significant otherwise, two-tailed t-test. **h**, Same as (**g**) but for changes in *L_max_*. **i**, Same as (**g**) but for changes in the ability to direct licks away from the midline, quantified using |*Θ_shoot_*|. **j**, Same as (**g**) but for changes in the variability of lick angles, quantified using the standard deviation of *Θ_shoot_*.

With the light powers we used (4 mW each hemisphere; Methods), light within 1 mm distance reduced mean spike rate of putative pyramidal cells (Extended Data Fig. 2a-c) by 91%, light at ∼1.5 mm away by 61%, and ∼3 mm away by 19% in behaving animals (Extended Data Fig. 2d). Interestingly, the mean spike rate of putative fast spiking (FS) neurons at ∼3 mm away was also reduced by 19%, rather than showing an increase due to photoactivation, suggesting that the decreased activity of both pyramidal and FS neurons was likely due to a reduction of cortical input. In contrast, light shined within 1 mm increased the mean spike rate of FS neurons by 739% and at ∼1.5mm by 140%.

#### S1TJ is required for proper targeting

Somatosensory inputs not only provide information about external objects but also enable the proprioceptive sensing of the body’s position in space ^24^ for motor control ^25^. Missing sensory feedback can render seemingly effortless manipulations surprisingly difficult even though motor capability per se is unchanged ^26^. Inhibiting S1TJ randomized the targeting angle of licks (Movie 3), shown by the increased standard deviation (SD) in *Θ* of individual licks (Fig. 3d; shadings around *Θ* traces) and the SD in *Θ_shoot_* at given time points for individual mice (Fig. 3j and Extended Data Fig. 2j). Despite the increased variability in targeting, the ability to direct licks away from the midline (i.e. |*Θ_shoot_* - 0°| or simply |*Θ_shoot_*|) was uncompromised (Fig. 3i). Inhibiting S1TJ also did not shorten the length of licks (Fig. 3h). Taken together, this suggests that S1TJ inhibition left intact the core motor capabilities required for tongue protrusions and licking, but corrupted these commands and their proper targeting, possibly due to missing sensory feedback.

In contrast, when inhibiting ALM (Fig. 3c), mice not only had difficulty directing licks away from the midline (Fig. 3i, Extended Data Fig. 2i), but also showed decreased length of lick (Fig. 3h, Extended Data Fig. 2h). The variability of licks, however, did not change (Fig. 3j, Extended Data Fig. 2j). Inhibiting M1B (Extended Data Fig. 2e) only caused a moderate increase in the variability of lick angle and no decrease in lick length. Inhibiting S1BF (Fig. 3e, Extended Data Fig. 2g-j) and S1Tr (Extended Data Fig. 2f) did not change any aspects of lick control (Fig. 3g-j).

#### ALM is critical for sequence initiation

ALM has been shown to be important in motor preparation of directed single licks to obtain water reward ^18, 20^. Here, we found that inhibiting ALM at sequence initiation strongly suppressed the production of licking sequences (Fig. 3c; Movie 4). In 3 of the 5 mice, licks were largely absent (Fig. 3g and Extended Data Fig. 2g). Inhibiting S1TJ caused more moderate suppression (Fig. 3d), and there was no obvious change when inhibiting other regions (Fig. 3e,g and Extended Data Fig. 2e-g). When applied at mid-sequence, ALM inhibition also suppressed the production of licks, although less strongly. Inhibiting any other region at mid-sequence showed little effect (Fig. 3 and Extended Data Fig. 2).

#### Anterior cortex activity is collectively required for sequence termination

Previous studies have attributed M1B almost exclusively to body and limb control. To our surprise, when inhibiting M1B at water consumption, mice were impaired at stopping ongoing sequences (Extended Data Fig. 2e, right; Movie 5). This prolonged licking was not due to additional attempts to reach the port for water as mice continuously made successful contacts. However, we also observed a trend toward an increase of lick production when inhibiting S1TJ, and more subtly, S1BF and S1Tr (Fig. 3 and Extended Fig. 2).

To test the possibility that inhibition of multiple areas caused persistent lick bouts due simply to spread of inhibition into M1B, we repeated all the above experiments with half of the illumination power (2 mW). The effects of ALM inhibition on sequence initiation, tongue length and angle control, and of S1TJ inhibition on angle control, remained consistent with, though much weaker than, our previous results using higher power (Extended Data Fig. 2k-n). At consumption, however, inhibiting M1B resulted in similar deficit in terminating ongoing sequences. Surprisingly, this paradoxical effect also became evident when inhibiting ALM and S1TJ (Fig. 3f). Therefore, our observation that inhibition of multiple anterior cortical areas produces a deficit in sequence termination was not due to spread of inhibition into M1B.

Rather, our results indicate that sequence termination is an active process mediated collectively by multiple regions in the anterior cortex. Since our data show that ALM and S1TJ played active roles in tongue control, inhibiting them with high power presumably impaired both sequence execution and termination. Low-power inhibition appeared to spare the control of execution more than that of termination, revealing deficits in the latter.

#### M1TJ inhibition impairs motor control of licking

Functional mapping and anatomical tracing from previous studies have shown that M1TJ is involved in the control of tongue/jaw motions ^13, 18^ and that it communicates with S1TJ ^13^. To test the role of M1TJ in sequence control, we performed additional inhibition experiments in a separate group of mice. M1TJ inhibition showed similar effects as ALM inhibition, including suppressed lick production, shortened tongue length and reduced angle modulation (Extended Data Fig. 2o,p).

### Single-unit responses tile sequence progression

We used silicon probes to record from multiple brain regions from both hemispheres (Fig. 4a) during the task and obtained a total of 1312 single-units and 284 multi-units (Methods; Extended Data Fig. 3a-e) from 51 recording sessions. Perievent time histograms (PETHs) of single-unit spiking (Fig. 4b,d,f) exhibited a wide variety of patterns prior, during, and after sequence execution.

**Fig. 4.**
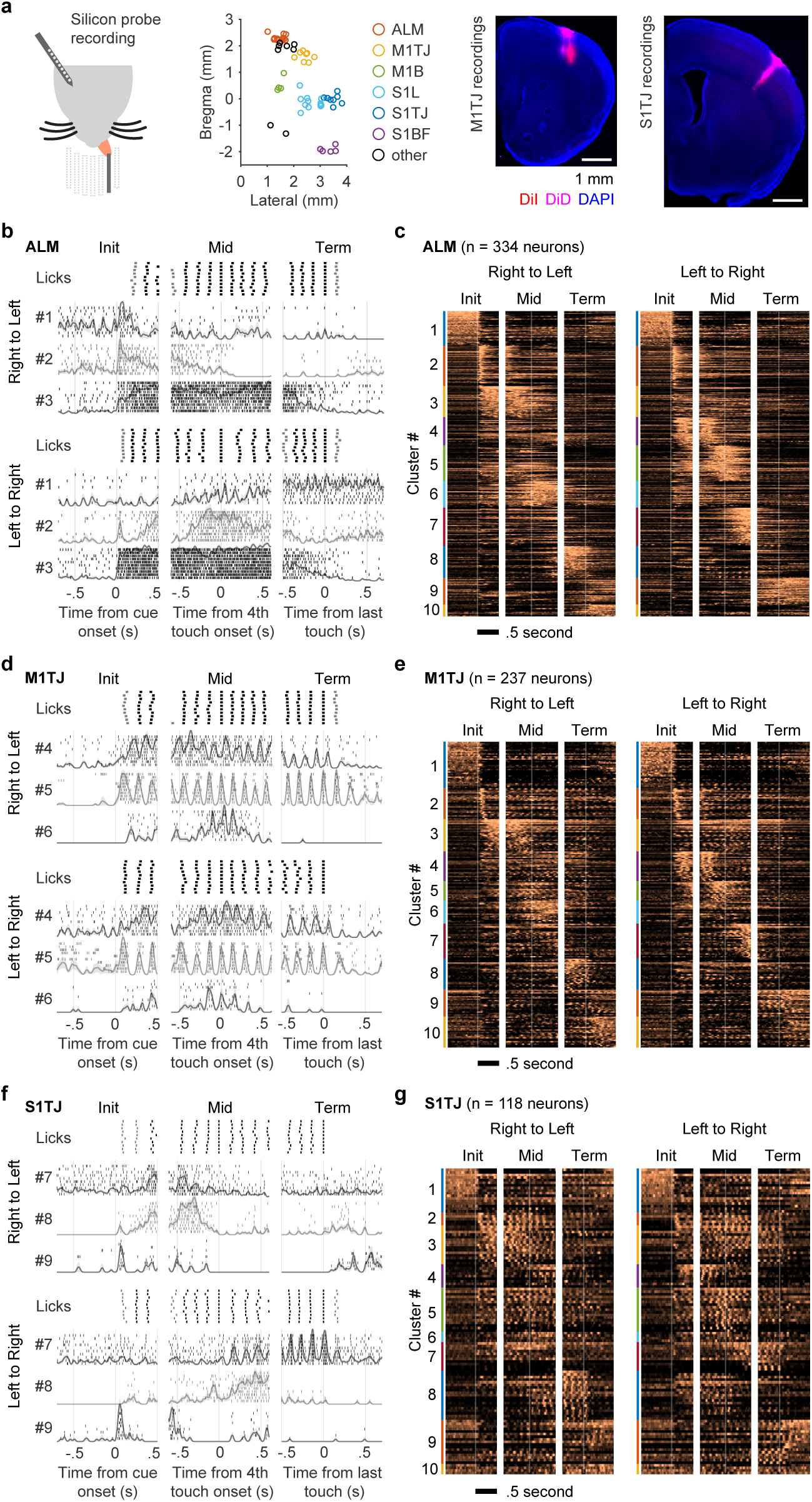
Single-unit activity tiles sequence execution. **a**, Left, silicon probe recording during the sequence licking task. Middle, histologically verified locations of silicon probe recordings. Recordings were made in both hemispheres but are illustrated for one. The plotted coordinates were randomly jittered by ± 0.05 mm to avoid visual overlap. Right, example sections showing the dye-labeled probe tracks. DiI and DiD were used in different penetrations. **b**, Responses of three simultaneously recorded ALM neurons aligned with right to left (top) or left to right (bottom) licking sequences at initiation (left), mid-sequence (middle) and termination (right; defined as the onset of the last consummatory lick). For each sequence direction, the first row shows rasters of lick times (touches in black and misses in gray) from 10 selected trials (Methods). Stacked below are spike rasters and the corresponding PETHs from the same 10 trials for each example neuron. **c**, Normalized PETHs of all ALM neurons plotted as heatmaps. Neurons are grouped together by functional cluster (Results) and labeled by color bands. Time zero for each period is marked by the vertical lines and is consistent with (**b**). **d**, Same as (**b**) but for three example neurons from M1TJ. **e**, Same as (**c**) but for all M1TJ neurons. **f**, Same as (**b**) but for three example neurons from S1TJ. Neurons #7 and #8 were both tuned to licking towards the right. In addition, neuron #7 fired more strongly when the tongue contacted the port. **g**, Same as (**c**) but for all S1TJ neurons.

To discover the main themes of single neuron level encoding in an unsupervised way, we pooled neurons from all brain regions and clustered them based on their PETHs using non-negative matrix factorization (NNMF, a technique closely related to principal component analysis, or PCA, and K-means clustering; Methods). We chose NNMF for its simplicity as a linear technique, its lack of an orthogonality constraint (present in PCA), and its duality as both a dimensionality reduction and clustering method.

We computed 10 clusters of PETHs (Fig. 4c,e,g and Extended Data Fig. 3f). Cluster #1 neurons showed high activity outside sequence execution. Cluster #2 showed transient activation at sequence initiation. In contrast to the first two, all other clusters exhibited directional selectivity at specific stages, temporally tiling the behavioral sequences. Interestingly, neurons in clusters #9 and #10 reached peak activation when licking sequences stopped. Our finding of recurring PETH motifs was not due to our use of the specific NNMF method or the chosen number of clusters, as a different method yielded consistent results (Methods; Extended Data Fig. 4g-j). Across NNMF clusters, activity of many neurons was smoothly modulated over hundreds of milliseconds; others, especially common in M1TJ and S1TJ, showed rapid modulation with individual licks. None of the clusters was biased to represent a small minority of neurons (Extended Data Fig. 3k). However, the different cortical regions harbored different proportions of the 10 clusters. In contrast to ALM, M1TJ and S1TJ, the 7 clusters from #2 through #8 made up only ∼1/3 of all S1BF neurons (Extended Data Fig. 4l). With neurons pooled from all regions, different cortical depths contained similar proportions of clusters (Extended Data Fig. 3m). Although the cluster proportions in individual regions could differ by cortical depth (e.g. cluster #2 units appeared less in S1TJ than in ALM or M1TJ), it is unclear whether or not this was due to an insufficient number of neurons once subdivided by depths and clusters (Extended Data Fig. 3n).

In sum, we observed both variety and commonality at the level of single-neuron responses. However, do patterns of activity arising from these single-unit responses encode behavioral variables important for sequence control?

### Hierarchical population coding of behavioral variables across cortical areas

In our sequence licking task, the brain needs to encode instantaneous tongue length (*L*) and angle (*θ*), presumably both for motor output and sensory feedback. The encoding of velocity (*L’*) could also be used to indirectly control tongue position. However, encoding only instantaneous kinematics is not enough, as mice need to string individual licks into sequences. Sequences in one direction also require a different organization of licks compared with sequences in the other direction. One way to organize a sequence is to have a slowly varying encoding of expected target position, regardless of the presence or absence of a lick at a given moment in time. Another, mutually non-exclusive, way is to separately encode sequence direction (*D*) and relative sequence time (*τ*). The variable *τ* can also serve as a proxy for sequence progress or “distance to goal”. The five behavioral variables, *L*, *L’*, *θ*, *D* and *τ*, were all recorded (or derived) with a temporal resolution of 2.5 ms (Fig. 5a). Conveniently, any pair of these variables is uncorrelated (Extended Data Fig. 4a). Therefore, being able to encode one is of little or no help to encode any other.

**Fig. 5.**
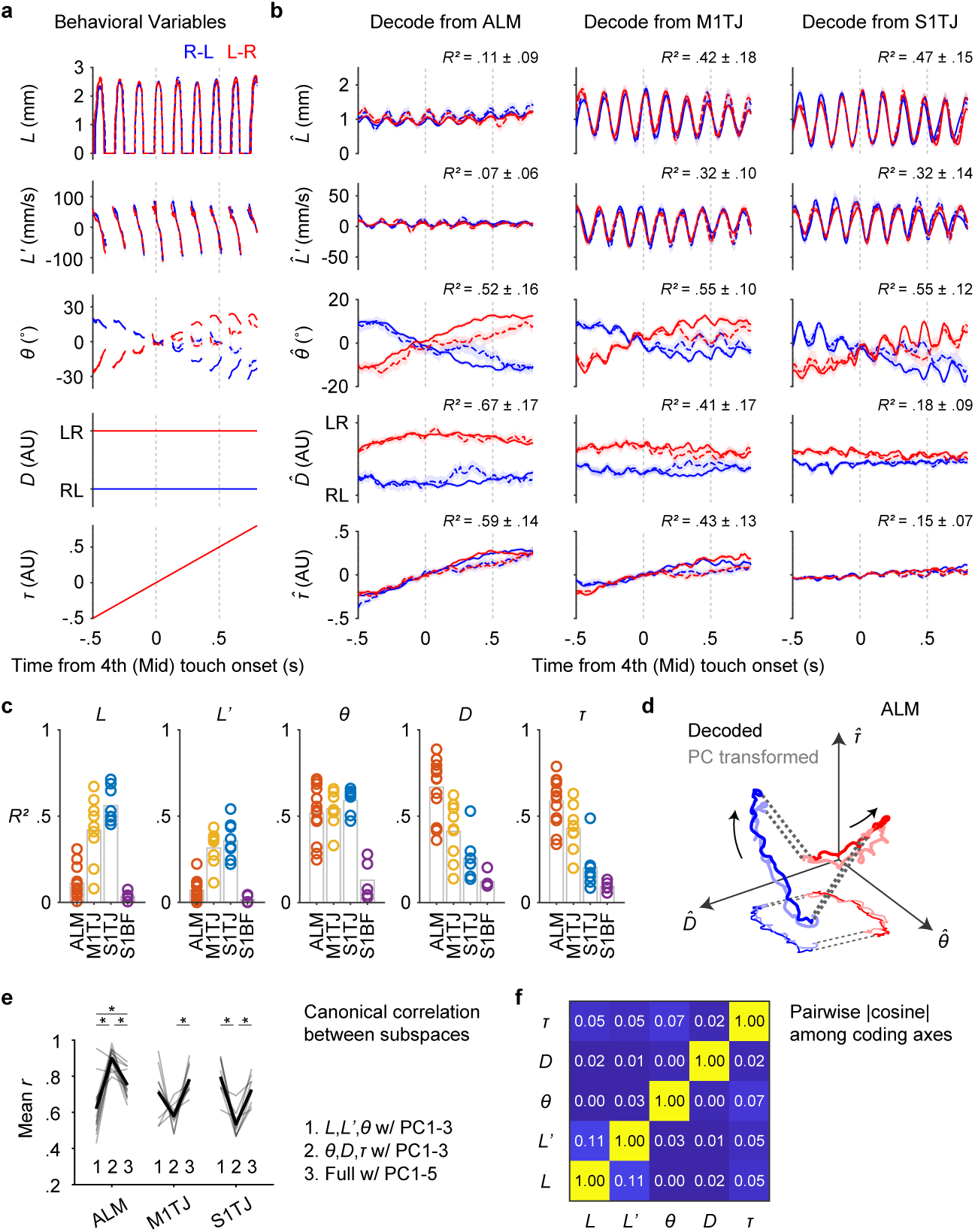
Populations code behavioral variables at increasing levels of abstraction across cortical areas. **a**, The five recorded (or derived) behavioral variables averaged across trials (n = 2684 trials; mean ± 99% bootstrap confidence interval) with standard sequences (solid lines) or backtracking sequences (dashed lines), each of which is either right to left (blue) or left to right (red). Time along the x-axis is aligned to the fourth lick in the sequence. Time points where more than 80% of trials did not have observations are not plotted. **b**, Decoding of the five behavioral variables (rows) from populations recorded in ALM, M1TJ, and S1TJ (columns). Cross-validated *R*^2^ for each region and variable is given (mean ± SD; ALM, n = 13 recordings; M1TJ, n = 9 recordings; S1TJ, n = 8 recordings). Same plotting conventions as in (**a**). **c**, Goodness of fit for linear models that predict each of the five behavioral variables, quantified by cross-validated *R*^2^. Each plot symbol shows one recording session. **d**, Neural trajectories from ALM (mean; n = 13 recordings) during standard right to left (blue) and left to right (red) sequences (linked by dashed lines). Arrows indicate the direction of time. The decoded trajectories (darker thick curves) are overlaid with the trajectories (lighter thick curves) in the space of the top 3 PCs but linearly transformed to align with the decoded ones. A projection of these trajectories is shown in the *D*-*θ* plane with thinner and lighter curves. **e**, Mean canonical correlation coefficients (*r*) of each neural population (gray trace) across three conditions. The average mean *r* values for each condition are shown in black (ALM, n = 13 recordings; M1TJ, n = 9 recordings; S1TJ, n = 8 recordings). ∗ p < 0.002, not significant otherwise, paired two-tailed permutation test. **f**, Absolute pairwise cosine values among coding axes (mean; n = 35 recordings).

Linear models, though simple, are powerful methods to uncover information encoded by a population of neurons ^27^. For each recording session, we performed separate linear regressions (Methods) to obtain unit weights (and a constant) for each of the five behavioral variables, such that a weighted sum of spike rates from simultaneously recorded units (31 ± 14 units; mean ± SD) plus the constant best predicted the value of a behavioral variable. We used cross-validated *R*^2^ values to quantify how well the recorded population of neurons encoded each behavioral variable. Importantly, we used only data from standard sequences to fit the models, but used these same models to decode both standard and backtracking sequences. The five behavioral variables could be decoded from population activity even for single trials (examples from ALM and S1TJ are shown in Extended Data Fig. 4b,c).

Despite our observation that ALM inhibition uniquely produced a deficit in maintaining *L* during sequences (Fig. 3), neuronal populations in this area showed weak encoding of instantaneous *L* and *L’* (Fig. 5b,c). In contrast, the encoding was much stronger in M1TJ and S1TJ. This suggests that ALM may control *L* indirectly over longer timescales via a signal to other brain regions, or is simply permissive for control of *L*. We consider the latter possibility unlikely because stimulation of ALM drives licking ^11, 18, 28^. Our previous results showed that the pattern of *L* and *L’* of individual licks are the same regardless of sequence direction or whether or not mice were relocating a port during backtracking sequences (Fig. 2d). The overlapping decoded *L* and *L’* traces for M1TJ and S1TJ are consistent with this behavioral invariance.

ALM, M1TJ and S1TJ, but not S1BF, all showed strong encoding of *θ* (Fig. 5b,c, Extended Data Fig. 4d). When detecting backtracking, mice licked back to a previous angle to relocate the port and then progressed through the rest of the sequence. The opposing deflections in the decoded *θ* from backtracking trials (Fig. 5b, dashed curves for *θ*) matched this behavior. The traces of decoded *θ* in M1TJ and S1TJ contained rhythmic fluctuations that were absent in ALM, despite similar overall levels of encoding of *θ* (*R*^2^ values). These fluctuations indicate that M1TJ and S1TJ encoded *θ* in a more instantaneous manner, whereas ALM encoded *θ* in a continuously modulated manner that may provide a control signal for the intended lick angle or represent the position of the target port.

Higher-level cortical regions are in part defined by the presence of more abstract (or latent) representations of sensory, motor and cognitive variables. Compared with *L*, *L’* and *θ*, which describe the kinematics of individual licks, sequence direction (*D*) and relative sequence time (*τ*) describe more abstract motor variables. In ALM we found the strongest encoding of both *D* and *τ* (Fig. 5b,c). Encoding of *D* and *τ* became progressively weaker in M1TJ, S1TJ, and S1BF, respectively. Overall, these results reveal a neural coding scheme with increasing levels of abstraction across S1TJ, M1TJ and ALM during the execution of flexible sensorimotor sequences.

### Coding of behavioral variables for sequence generation dominates cortical activity patterns

Good decoding may come from a small fraction of informative units rather than dominant activity patterns across a population. More generally, we wondered whether the coding axes (i.e. coefficient vectors) of the five behavioral variables captured the dominant activity in the neural space, or were only related to minor components leaving the main dynamics unexplained. To determine this requires comparing the similarity between activity patterns captured by the coding axes and the dominant patterns in population activity identified in an unsupervised manner. In each recording session, we obtained neural trajectories in the coding subspaces (the subspaces spanned by coding axes) via linear decoding and trajectories in principal component (PC) subspaces (the subspaces spanned by the first few PCs) via PCA. Trajectories in PC subspaces depict dominant patterns in population activity but the PCs per se need not have any behavioral relevance. To see if neural trajectories in the coding and the PC subspaces were the same except for a change (rotation and/or scaling) in reference frame, we used canonical correlation (Methods) to find the linear transformation of the two trajectories such that they were maximally correlated.

After transformation, trajectories of the ALM population in the subspace of the top three PCs very well aligned (Fig. 5d) and correlated (Fig. 5e; group 2 in ALM) with the trajectories in the subspace encoding *θ*, *D,* and *τ*. This indicates that the most dominant neural activity patterns in the ALM population in fact encoded *θ*, *D* and *τ*. Since ALM minimally encoded *L* and *L’*, including these to the coding subspaces decreased the correlation with PC trajectories (Fig. 5e; group 1 and 3 in ALM). The decoded trajectories and PC trajectories in M1TJ and S1TJ also showed strong correlation but only when the coding subspaces included *L* and *L’*.

Across regions, the sum of variance explained (VE) by the five coding axes reached about half of that by the top five PCs (Methods; Extended Data Fig. 4e). The five coding axes were largely orthogonal among each other (Fig. 5f). This indicates that they not only captured dominant neural dynamics but also did so efficiently with little redundancy.

### Reward modulates dominant activity patterns in ALM

The coding axis of *τ* was identified by fitting models between neural activity and relative sequence time. However, if *τ* faithfully represents time, the downward deflection of traces from backtracking sequences (Fig. 5b) should not appear, as time goes on regardless of what the animals do. Therefore, the patterns suggest a representation of a “distance to goal.” In the context of the motor sequence, does the goal represent the initial arrival at the last port position, or the delivery of the water droplet, or finishing water consumption, etc?

In ALM, we found single neurons (Fig. 6a) that fired actively during sequence execution but abruptly decreased their firing at the time when the tongue touched water droplets. Mice continued to emit ∼5 consummatory licks (Fig. 6b) with similar, if not more strongly modulated, kinematics and lick force (Fig. 6c). We decoded behavioral variables around the consumption period using linear models fitted during sequence execution (i.e. without data from the consumption period). The *τ* decoded from ALM populations (Fig. 6d; top) immediately decreased around the first contact (∼0 s) with water and diminished (∼0.5 s) before a noticeable decrease in lick production. This suggests that *τ* carried a reward expectation signal that smoothly increased as mice approached water delivery regardless of sequence direction or lick angle, but was suppressed by a delay of progress when backtracking occurred, and terminated at the time when the tongue detected water, despite continued licking movements. Curiously, the *D* coding (Fig. 6d; middle) followed a similar time course as *τ*, although the implication of an interaction between sequence direction and reward is unclear.

**Fig. 6.**
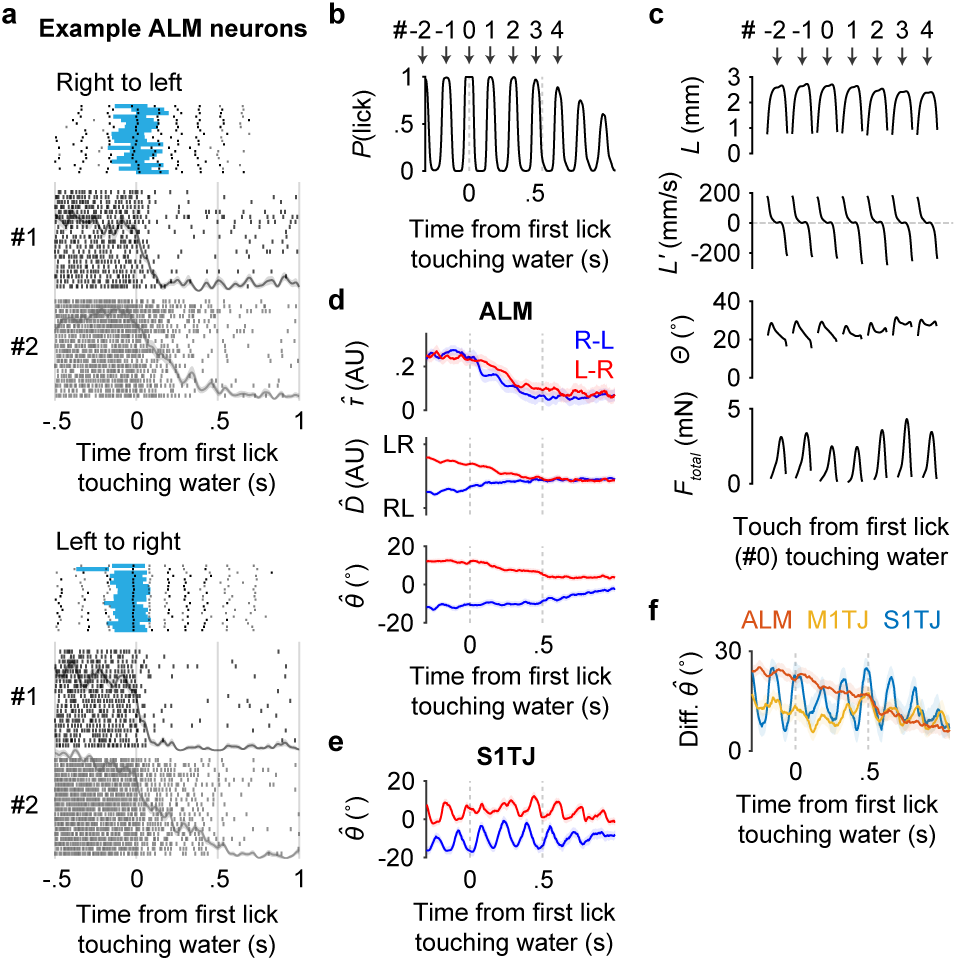
Reward modulation of activity in ALM. **a**, Responses of two simultaneously recorded ALM neurons (#1 and #2) aligned at the first lick (specifically the middle of a tongue-out period) that touched water reward. For each sequence direction, shown at top are rasters of lick times (touches in black and misses in gray) and the duration of water delivery (blue) from 20 selected trials (Methods). Stacked below are spike rasters and the corresponding PETHs from the same 20 trials for each example neuron. **b**, The probability of licking (i.e. tongue-out) as a function of time. Licks are sequentially indexed with respect to the first lick (#0) touching the water. **c**, Patterns of kinematics and force for single licks around the first lick (#0) touching water (n = 25289 trials; mean ± 95% bootstrap confidence interval). The duration of individual licks was normalized. The total force (*F_total_*) is the vector sum of vertical and lateral forces. **d**, Decoding of *τ*, *D* and *θ* from neuronal populations recorded in ALM (mean ± 99% bootstrap confidence interval; n = 13 recordings) in right to left (blue) or left to right (red) trials around the consumption period. **e**, Similar to (**d**) but decoding *θ* from S1TJ (n = 8 recordings). **f**, The difference between the decoded *θ* traces in right to left versus left to right trials (mean ± 99% bootstrap confidence interval; ALM, n = 13 recordings; M1TJ, n = 9 recordings; S1TJ, n = 8 recordings).

The *θ* coding in ALM during the consumption period exhibited a more complex time course (Fig. 6d, bottom). A moderate reduction in the separation of decoded *θ* in the two sequence directions occurred once the tongue touched the water. However, it remained separated toward the end of the lick bout. In contrast, the overall separation in *θ* coding from S1TJ (Fig. 6e,f) and M1TJ (Fig. 6f) was not altered by the detection of water, and the amplitude of rhythmic fluctuations was consistent with individual patterns of licking force (compare S1TJ and M1TJ traces in Fig. 6f with the *F_total_* in Fig. 6c). Overall, our results show that reward-related modulation is a prevalent feature in ALM but not S1TJ or M1TJ coding.

### ALM encodes upcoming sequences

Sequences with opposite directions were performed alternatively across trials (Extended Data Fig. 5a). Mice performed the task in darkness. Hairs and whiskers around the mouth were trimmed short to avoid contact with the port. Therefore, there was no external cue indicating which side a mouse should start a sequence. Nevertheless, expert mice were usually able to initiate sequences from the correct side without exploring the other (Extended Data Fig. 5b). This suggests that the information about target position (*TP*) was maintained internally during ITIs. Brain regions maintaining such information may be important in organizing individual sequences together to form a higher level sequence.

In ALM recordings, we found simultaneously recorded units that fired persistently to specific *TP* during ITI (Extended Data Fig. 5c). A linear model fitted using data within 1 s before the cue onset showed smooth population decoding of *TP* across the span of many trials (Extended Data Fig. 5d). On average, ALM populations showed stronger encoding of *TP* (Extended Data Fig. 5e,f) compared to other regions, although overall it was weaker than either the encoding of *D* or *θ* during sequence execution. Interestingly, none of the regions, including ALM, encoded time or a distance to trial start (Extended Data Fig. 5g), perhaps because our ITI contained an exponential portion (Methods) whose flat hazard function made time to trial start unpredictable^10^.

At the beginning of a sequence, *D* and *θ* both matched the preceding *TP*. Given that *D* and *θ* were encoded independently (i.e. in orthogonal neuronal subspaces) (Fig. 5f), we wondered whether the coding of *TP* was more relevant to later *D* or *θ*, or neither. We used linear models fitted during the ITI to decode from neural activity during sequence execution. The resulting traces from two sequence directions crossed at the mid-sequence (Extended Data Fig. 5h). This revealed that *TP* coding reflects a persistent encoding of intended tongue angle in the absence of licks.

## Discussion

Rodents are highly tactile species and use rich orofacial behaviors to interact with the environment. We found that mice exhibited a remarkable ability to learn complex licking sequences. The sequence structure within a trial and the alternation of different sequences allowed investigation of additional dimensions of motor control beyond simple cue-response contingencies or repetitive movement sequences. Mice were able to incorporate arbitrarily defined new transitions into the existing repertoire of learned sequences, and quickly decide whether, and how, to change an ongoing motor program based on tactile feedback. These feedback-driven alterations involved complete reversals of direction and reorganization of the motor sequence. Ongoing performance in our task, even in well-trained mice, required activity in motor cortex, although we speculate that our task also depends on basal ganglia and cerebellar circuits. In general, the mechanisms within motor cortex that allow sensory feedback to integrate with unfolding motor programs is a long-standing and active area of research ^29–33^.

To perform a sequence of directed licks, the nervous system must generate motor commands to lengthen the tongue and modulate its angle. Optogenetic inhibition of ALM impaired both the length of each lick and the ability to direct licks away from the midline. These results partly support recent work showing that inhibition of ALM impairs aspects of tongue kinematics in a cued-licking task ^34^. Electrophysiological recordings in ALM revealed strong encoding of *θ* but surprisingly not *L* or *L’* (which can indirectly produce *L*). This suggests that ALM may control *L* indirectly over longer timescales via a signal to other cortical or subcortical regions. In contrast to the rhythmically fluctuating *θ* trajectories decoded from M1TJ and S1TJ populations, ALM populations encoded an intended *θ* that varied smoothly over time. Our data do not rule out that ALM may also encode an intended *L*, or target distance. In our task mice used similar lick lengths across different port positions and we thus cannot distinguish signals for intended *L* versus task engagement. A task requiring mice to lick at different distances may help to resolve this question.

Precise motor outputs in skilled behaviors typically rely on sensory feedback ^25, 35^. Optogenetic inhibition of S1TJ produced distinct motor deficits from those observed with ALM inhibition. With S1TJ inhibition, mice could direct licks laterally to the normal extent and with normal length. However, lick angles became randomized and mice could no longer perform the correct sequences. Interestingly, neurons in S1TJ not only encoded *θ*, but also *L* and *L’*. This suggests that the system either does not need sensory feedback about tongue length to guide the licking sequence, presumably because the required lick lengths were always the same and could in principle be controlled in open-loop, or that the inhibited *L* and *L’* signals in S1TJ could be compensated by redundant information in other brain regions.

Performing different sequences in our task not only required precise control of single licks but also the appropriate sequencing of different licks. Our data reveal encoding and control of behavior at multiple levels of abstraction across cortical areas. Zebra finch birdsong, in which syllables are organized into sequences to form songs, has been a powerful model for dissecting neural control over motor sequences and has revealed a specialized circuit for hierarchical motor control ^36^. In primates, control of movements is distributed across multiple primary and premotor areas defined cytoarchitecturally and functionally. Medial premotor areas including the supplementary motor area (SMA) and pre-SMA play a prominent role in controlling the sequential organization of movements, while M1 activity may also be involved in sequence organization but in general is more closely related to imminent and ongoing movements ^37–40^. While different cortical areas show functional distinctions regarding the level of abstraction of behavioral variables encoded, multiple somatomotor cortical areas contain corticospinal neurons and thus have the potential to exert fairly direct control of muscles. In rodents, the organization of motor areas and their functional relationships are so far less clear. Our results show a functional organization of somatomotor areas that reflects encoding and control of task-related behavioral variables with increasing levels of abstraction, with S1TJ and M1TJ activity more tightly linked to instantaneous kinematics, and ALM to variables at the sequence level, specifically a smooth representation of intended lick angle (*θ*), sequence direction (*D*), and distance to reward (*τ*).

To uniquely represent a behavioral state at a moment within a given sequence, it is sufficient to either encode *θ* or encode both *D* and *τ*. Interestingly, the ALM populations encoded all three variables. This apparent redundancy may serve a computational role from a dynamical systems perspective on motor control ^41^. Each point in the state space of a smooth dynamical system is associated with a unique trajectory ^42^. Thus, if the system needs to produce two licking sequences, the two trajectories must be kept adequately separated so that they do not intersect in the presence of noise and thereby turn one sequence into the other ^43^. Encoding all three variables avoids trajectories crossing at the middle lick and maximizes the distance of separation (Fig. 5d). In addition, we found that *τ* exhibited properties of a reward expectation signal, and *D* and *θ* also showed reward-related modulations. This is consistent with recent work showing reward-related activity in ALM that depends on output from the cerebellar dentate nucleus ^44^. These variables may contribute to other processes other than sequence execution, such as representing instantaneous action values for motor learning.

Prior work showed that ALM holds working memory or motor preparation signals for upcoming lick direction in delayed response tasks, bridging the sensory-to-motor transformation ^6–12, 45^. Consistent with these results, we first found that optogenetic inhibition of ALM at cue onset strongly impaired the initiation of licking sequences. Before the go cue, ALM populations persistently encoded the upcoming sequence throughout the ITI. Furthermore, this signal is closely related to *θ*, but not *D*, suggesting that it represents the target position rather than a target sequence direction, despite the two being correlated at sequence initiation. Unlike previous tasks, our task does not have an explicit sensory event indicating the side on which to initiate a delayed sequence. However, the last port touch in a trial could be considered to serve as the “sensory event” and the location of the first lick in the next trial as the “response”. Thus, our findings generalize the forms of delayed sensory-to-motor transformation involving ALM and provide insights into how the brain solves a sequence of sequences.

Inability to stop an ongoing movement sequence can be as devastating as the inability to initiate one. When we optogenetically inhibited M1B at water consumption, mice showed difficulty in terminating an ongoing lick bout. This continued licking was not due to failure of the tongue to reach the port to retrieve water. It is also unlikely that inhibition prevented mice from sensing the water because: 1) cortical representations of water taste can be found in gustatory cortex ^46, 47^, which we did not inhibit; 2) inhibiting any region in the anterior dorsal cortex not just somatosensory cortex, induced a similar deficit in sequence termination; and 3) decreasing the stimulus light intensity by half in ALM and S1TJ resulted in stronger deficits, not weaker. These results suggest that multiple anterior cortices play a role in active termination of an ongoing sequence, perhaps via convergence of activity on a downstream target such as the basal ganglia circuits that are critical for both starting and stopping movement sequences ^16^.

Together, our results from behavior, population electrophysiology and optogenetics allowed us to define a functional network within mouse sensory and motor cortices that governs execution of flexible, feedback-driven sensorimotor sequences.

## Acknowledgements

We thank William Olson, Rajan Dasgupta, Yi-Ting Chang, Varun Chokshi, Jeremiah Cohen, Michael Economo, and Karel Svoboda for comments on the manuscript; Ali Aly, Yun Hwang, Hokin Deng for assistance with experiments and data curation; Varun Chokshi and William Olson for sharing animals; Bilal Bari and Jeremiah Cohen for VGAT-Cre;Ai32 mice; Travis Babola for suggestions on the hearing loss experiment. This work was supported by NIH grants R01NS089652 and 1R01NS104834-01 to D.H.O., and center grant P30NS050274.

## Author contributions

D.X., Y.C., A.M.D., N.C.H., M.D., and L.Z. performed experiments. D.X. developed custom software, hardware, analysis code and analyzed data, with input from all authors. D.X., N.C.H., M.D., and D.H.O. wrote the paper with input from all authors.

## Materials & Correspondence

Requests for materials and correspondence should be directed to D.H.O.

## Movie Legends

**Movie 1. Example performance in a standard sequence (related to Fig. 1)** Top, high-speed video capturing the bottom and side views of the mouse. The tracked base and the tip of the tongue are labeled by red asterisks. Bottom, time-aligned behavioral variables and events similar to Fig. 1d. A moving bar indicates the position of the current frame. The playback was slowed down 5-fold.

**Movie 2. Example performance in a backtracking sequence (related to Fig. 2)** Same conventions as for Movie 1.

**Movie 3. Example trial with inhibition in S1TJ at sequence initiation (related to Fig. 3)** Similar conventions as for Movie 1 but showing the waveform of optogenetic inhibition (instead of the vertical and lateral lick forces).

**Movie 4. Example trial with inhibition in ALM at sequence initiation (related to Fig. 3)** Same conventions as for Movie 3.

**Movie 5. Example trial with low-power inhibition in M1B at consumption (related to Extended Data Fig. 2)** Same conventions as for Movie 3.

**Extended Data Table 1. Mouse information**

## Methods

### Mice

All procedures were in accordance with protocols approved by the Johns Hopkins University Animal Care and Use Committee. Mice were kept in a reverse light-dark cycle with each phase lasting 12 hours. Prior to surgery, mice were housed in groups of up to 5, but afterwards housed individually. Ten mice (8 male, 2 female) were obtained by crossing VGAT-IRES-Cre (Jackson Labs: 028862; B6J.129S6(FVB)-Slc32a1^tm2(cre)Lowl^/MwarJ)^48^ with Ai32 (Jackson Labs: 012569; B6;129S-Gt(ROSA)26Sor^tm32(CAG–COP4*H134R/EYFP)Hze^/J) ^49^ lines. Two (1 male, 1 female) were heterozygous VGAT-ChR2-EYFP (Jackson Labs: 014548; B6.Cg-Tg(Slc32a1-COP4*H134R/EYFP)8Gfng/J) ^50^ mice. Eight (6 male, 2 female) were wild-type mice, including seven C57BL/6J (Jackson Labs: 000664) mice and one wild-type littermate of the VGAT-ChR2-EYFP mice. Two were male TH-Cre 1 (Jackson Labs: 008601; B6.Cg-7630403G23Rik^Tg(Th-cre)1Tmd^/J) ^51^ mice. Two (1 male, 1 female) were Advillin-Cre (Jackson Labs: 032536; B6.129P2-Avil^tm2(cre)Fawa^/J) ^59^ mice. Mice ranged in age from ∼2-9 months at the start of training. A set of behavioural testing sessions typically lasted ∼1 month.

### Surgery

Prior to behavioural testing, mice underwent the implantation of a metal headpost. For surgical procedures, mice were anesthetized with isoflurane (1-2%) and kept on a heating blanket (Harvard Apparatus). Lidocaine was used as a local analgesic and injected under the scalp at the start of surgery. Ketoprofen was injected intraperitoneally to reduce inflammation. All skin and periosteum above the dorsal surface of the skull was removed. The temporal muscle was detached from the lateral edges of the skull on either side and the bone ridge at the temporal-parietal junction was thinned using a dental drill to create a wider accessible region. Metabond (C & B Metabond) was used to cover the entirety of the skull surface in a thin layer, seal the skin at the edges, and cement the headpost onto the skull over the lambda suture.

To make the skull transparent, a thin layer of cyanoacrylate adhesive was then dropped over the entirety of the Metabond-coated skull and left to dry. A silicon elastomer (Kwik-Cast) was then applied over the surface to prevent deterioration of skull transparency prior to photostimulation. Buprenorphine was used as a post-operative analgesic and the mice were allowed to recover over 5-7 days following surgery with free access to water.

For silicon probe recording, a small craniotomy of about 600 μm in diameter was made for implantation of a ground screw. The skull was thinned using a dental drill until the remaining bone could be carefully removed with a tungsten needle and forceps. Following this, one or more craniotomies of about 1 mm in diameter were made over the sites of interest for silicon probe recording. Craniotomies were protected with a layer of silicon elastomer (Kwik-Cast) on top. Additional craniotomies were usually made in new locations after finishing recordings in previous ones.

### Task control

Task control was implemented with an Arduino-based system (Teensy 3.2 and Teensyduino), including the generation of audio (Teensy Audio Shield). Custom MATLAB-based software with a graphical user interface was developed to log task events and change task parameters. Touches between the tongue and the port were registered by a conductive lick detector (Svoboda lab, HHMI Janelia Research Campus), where the mouse acted as a mechanical switch that opened (no touch) or closed (with touch) the circuit. Any mechanical switch has electrical bouncing issues when a contact is weak and unstable. To handle bouncing during loose touches, we merged any contact signals with intervals less than 60 ms.

The auditory cue that signaled the beginning of each trial was a 0.1 s long, 65 dB SPL, 15 kHz pure tone. Touches that occurred during the auditory cue were not used to trigger port movement as they were likely due to impulsive licking rather than a reaction to the cue.

The lick port was motorized in the horizontal plane by two perpendicular linear stages (LSM050B-T4 and LSM025B-T4, Zaber Technologies), one for anterior and posterior movement and the other for left and right. A manual linear stage (MT1/M, Thorlabs) installed in the vertical direction controlled the height of the lick port. The motors were driven by a controller (X-MCB2, Zaber Technologies) which was in turn commanded by the Teensy board via serial interface communication. Although the linear stages were set up in cartesian coordinates, we specified the movement of the port using a polar coordinate system. For a chosen origin of the polar coordinates, the seven port positions were arranged in an arc symmetrical to the midline with equal spacing (in arc length) between adjacent positions (Fig. 1a). A movement of the lick port was triggered by the onset of a touch during sequence performance. A second port movement could not be triggered within 80 ms, which prevented mice from driving a sequence by constantly holding the tongue on the port (although we never observed such behavior). When a movement was triggered, the port first accelerated (477 or 715 mm/s^2^) until the maximal velocity (39.3 mm/s) was reached, then maintained the maximal velocity, and decelerated until it stopped at the end position. The acceleration and deceleration phases were always symmetrical, such that the maximal velocity might not be reached if the distance of travel was short. The movement was typically in a straight line. For 4 of the 9 mice, when the two positions were not adjacent (e.g. at backtracking and the following transition), the port would move in an outward half circle whose diameter was the linear distance separating the two positions. This arc motion minimized the chance of mice occasionally catching the port prematurely before the port stopped. Nevertheless, catching the port prematurely did not trigger the next transition in a sequence because, in this case, the port movement could only be triggered again after 200 ms from the start of backtracking (and 300 ms after the following touch). As a result, mice always needed to touch the port at the fully backtracked position in order to continue progress in a sequence.

Mice performed the task in darkness with no visual cues about the position of the port. To prevent mice from using sounds emitted by the motor to guide their behavior, we played two types of noise throughout a session. The first was a constant white noise (cutoff at 40 kHz; 80 dB SPL) and the second was a random playback (with 150-300 ms interval) of previously recorded motor sounds during 12 different transitions.

### Two-axis optical force sensors

A stainless steel lick tube was fixed on one end to form a cantilever. Mice licked the other free end, producing a small displacement (< ∼0.1 mm at the tip for 5 mN) of the tube. Two photointerrupters (GP1S094HCZ0F, Sharp) placed along the tube (Extended Data Fig. 1c,d) were used to convert the vertical and horizontal components of displacement into voltage signals. Specifically, the cantilever normally blocked about half of the light passing through, outputting a voltage value in the middle of the measurement range. Pushing the tip down caused the cantilever to block more light at the vertical sensor and thereby decreased the output voltage; conversely, less force applied at the tip resulted in increased voltage. For the horizontal sensor, pushing the tube to the left or right decreased or increased the voltage output, respectively. Output was amplified by an op-amp then recorded via an RHD2000 Recording System (Intan Technologies).

By design (the circuit diagram and the displacement-response curve are available in the GP1S094HCZ0F datasheet), the force applied at the tip of the lick tube and the sensor’s output voltage follow a near linear relationship within a range of forces. To find this range, we measured the voltages (relative to baseline) with different weights added to the tip. Excellent linearity (*R*^2^ = 0.9999) was achieved up to >20 mN (Extended Data Fig. 1d). In contrast, the maximal force of a lick was on average about 4 mN (Fig. 1f).

The motorization of the lick tube introduced mechanical noise to the force signals. The spectral components of these noises were mainly at 300 Hz and its higher harmonics, presumably due to the resonance frequency of the tube, whereas the force signal induced by licking occupied much lower frequencies. Therefore, we low-pass (at 100 Hz) filtered the original signal (sampled at 30 kHz/s) to remove the motor noise. Additional interference came from the 850 nm illumination light used for high-speed video, which leaked into the optical sensors (mainly in early experiments with 2 mice) and caused slow fluctuations in the baseline over seconds. To mitigate this slow drift, we used a baseline estimated separately for each individual lick as follows. We first masked out the parts of the signal when the tongue was touching the port, then linearly interpolated to fill in these masked out lick portions using the neighboring (i.e. no touch) values. These interpolated time series served as the baseline for each lick. Since the lick force was only a function of voltage change compared to baseline, the above procedure would at most negligibly affect the force estimation. Due to the dependency of this procedure on complete touch detection, we excluded 8 sessions from behavioral quantifications in Figs. 1 and 2 where only touch onsets were correctly registered.

### High-speed videography and tongue tracking

High-speed video (400 Hz, 0.6 ms exposure time, 32 µm/pixel, 800 pixels x 320 pixels) providing side- and bottom-views of the mouth region was acquired using a 0.25X telecentric lens (55-349, Edmund Optics), a PhotonFocus DR1-D1312-200-G2-8 camera, and Streampix 7 software (Norpix). Illumination was via an 850 nm LED (LED850-66-60, Roithner Laser) passed through a condenser lens (Thorlabs).

Three deep convolutional neural networks were developed (MATLAB, Deep Learning Toolbox) to extract tongue kinematics and shape from these videos. The first network classified each frame as “tongue-out”, if a tongue was present, or “tongue-in” otherwise. This network was based on a pretrained network, ResNet-50 ^52^, but the final layers were redefined to classify the two categories. A total of 37658 frames were manually labeled in which 1611 frames were set aside as testing data. Image augmentation was performed to expand the training dataset. A standard training scheme was used with a mini-batch size of 32 and a learning rate of 1×10^-4^ to 1×10^-5^. The fully trained network achieved a high accuracy in classifying the validation data (Extended Data Fig. 1a).

The second network assigned a vector from the base to the tip of the tongue in each frame classified as “tongue-out”. *L* and *θ* were derived from this vector (Fig. 1c). A total of 12095 frames were manually labeled in which 643 frames were used only for testing. The architecture and training parameters of this network are similar to those of the classification network except that the final layers were redefined to output the x and y image coordinates of the base, tip and two bottom corners (not used in analysis) of the tongue with mean absolute error loss. The regression error of the fully trained network in testing data was 3.1 ± 5.4° for *θ* and 0.00 ± 0.13 mm for *L* (mean ± SD). This performance was comparable to human level (Extended Data Fig. 1b). Specifically, a subset of frames (separate from testing data) were labeled by each of the five human labelers. The variability in human judgement was quantified by the differences between *L* and *θ* from individual humans and the human mean for each frame. We also computed the differences between *L* and *θ* from the network and the human mean for each frame. The two distributions showed a comparable variability, although the network showed small biases (*L*: humans 0 ± 0.11 mm, network −0.05 ± 0.10 mm; *θ*: humans 0 ± 5.7° SD, network 3.3 ± 5.5° SD; mean ± SD).

In a subset of trials and in frames classified as “tongue-out”, the third network, a VGG13-based SegNet ^53^, extracted the shape of the tongue by semantic image segmentation, i.e. classifying each pixel as belonging to a tongue or not. Human labelers used a 10-vertex polygon to encompass the area of the tongue in a total of 3856 frames. The training parameters were similar to the other networks except for a mini-batch size of 8 and a learning rate of 1×10^-3^.

### Behavioral training

Behavioural sessions occurred once per day during the dark phase and lasted for approximately an hour or until the mouse stopped performing, whichever came earlier. Mice would receive all of their water from these sessions, unless it was necessary to supply additional water to maintain a stable body weight. The amount of water consumed during behaviour was measured by subtracting the pre-session volume of water in the dispenser from the post-session volume. On days where their behaviour was not tested they received 1 ml of water. Mice were water restricted (1 ml/day) for at least 7 days prior to beginning training. Whiskers and hairs around the mouth were trimmed frequently to avoid contact with the port.

The precise position of the implanted headpost varied across mice, so each mouse required an initial setup of the lick port’s positions. The lick port moved in an arc with respect to a chosen origin (see Task Control). The origin was initially set at the midline of the animal and 2 mm posterior from the posterior face of the upper incisors. If there was any yaw of the head, the whole arc was rotationally shifted accordingly. The lick port’s z-axis was manually adjusted until the lick port was approximately 1 mm below the interface between upper and lower lips when the mouth was closed.

In initial training sessions, the distance between the leftmost (L3) and the rightmost (R3) lick port position was reduced, the radius of the arc was shortened, and the water reward was larger. As mice learned the task, both the L3 to R3 distance and the radius of the arc were gradually increased over a few days of training (Extended Data Fig. 1h). The difficulty of the task was increased whenever the mouse showed improvements in performing the task at a given set of parameters. The difficulty remained constant in two conditions: either when the maximum set of parameters had been met (a radius of 5 mm for males and 4.5 mm for females) or if the mouse appeared demotivated (typically indicated by a significant decrease in the number of trials and licks). During the initial training sessions, water was occasionally supplemented at other points during the sequence to encourage licking behaviour. The amount of water reward per trial was eventually lowered to ∼3 μL. For 3 of the 24 mice included in this study, we first trained them to lick in response to the auditory cue with the lick port staying at fixed positions. After mice responded consistently to the go cue, we shifted to the complete task with gradually increased difficulty. Although the 3 mice performed similarly to others when well trained, this procedure proved to be less efficient than beginning with the complete task.

Once a mouse had become adept at standard sequences, they were trained on the backtracking sequences. The first 9 fully trained mice were used in backtracking related analyses; later mice used for other purposes were not always fully trained in backtracking. For 5 of the 9 mice, we first trained them with backtracking trials in only one direction and added the other direction once they mastered the first. For 3 of the 9 mice, backtracking trials and standard trials were organized into separate blocks of 30 trials each. In developing this novel task, we tested subtle variations in the detailed organization of trial types, such as varying the percentage of backtracking trials in a block, or different forms of jumps in the port position. Details appear in Extended Data Table 1. Two of these 3 mice continued to performed the block-based backtracking trials during recording sessions. All 9 mice eventually learned backtracking sequences but showed mixed learning curves (Extended Data Fig. 1i,j). About 3 mice were more biased to previously learned standard sequences and tended to miss the port many times before relocating the lick port through exploration. The other 6 mice more readily made changes.

### Hearing loss

Hearing loss experiments were performed to exclude the possibility that mice used sounds produced by the motors to localize the motion of the lick port during sequence performance. To induce temporary hearing loss (∼27.5 dB attenuation) ^54^, we inserted two earplugs made of malleable putty (BlueStik Adhesive Putty, DAP Products Inc.) into the ear canal openings bilaterally under microscopic guidance. Earplugs were shaped like balls and then formed appropriately to cover the unique curvature of each ear canal. When necessary, the positioning of the earplugs was readjusted, or larger balls were inserted. Five well trained mice performed one “earplug” session and one control session. Mice did not have experience with earplugs prior to the earplug session. In earplug sessions, mice were first anesthetized under isoflurane to implant earplugs (taking 11-12.5 mins), then were put back to the homecage to recover from anesthesia (taking 10-11.5 mins), and performed the task after recovery. In control sessions, mice were anesthetized for the same duration (and to remove earplugs if necessary), and allowed to recover for the same duration before performing the task.

### Odor masking

Odor masking experiments were performed to exclude the possibility that mice used potential odors emanating from the lick port to localize its position during sequence performance. A fresh air outlet (1.59 mm in diameter) was placed in front of the mouse and aimed at the nose from ∼2 cm away with ∼45° downward angle. We checked the coverage of air flow (2 LPM) by testing whether a water droplet (∼3 μL) would vigorously wobble in the flow at various locations, and confirmed that both the nose and all the seven port positions were covered. Prior to the test session, head-fixed mice were habituated to occasional air flows when they were not performing sequences. In the test session, the air flow was turned off first and turned on continuously after the 100th trial (in four mice) until the end of the session, or turned on first and turned off after the 100th trial (in two mice). The air-off period served as the control condition for air-on period.

### Electrophysiology

Two types of silicon probe were used to record extracellular potentials. One (H3, Cambridge Neurotech) had a single shank with 64 electrodes evenly spaced at 20 µm intervals. The other (H2, Cambridge Neurotech) had two shanks separated by 250 µm, where each shank had 32 electrodes evenly spaced with 25 µm intervals. Before each insertion, the tips of the silicon probe were dipped in either DiI (saturated) or DiD (5-10 mg/mL) ethanol solution and allowed to dry. Probe insertions were either vertical or at 40° from the vertical line depending on the anatomy of the recorded region and surgical accessibility. Once fully inserted, the brain was covered with a layer of 1.5% agarose and ACSF, and was left to settle for ∼10 minutes prior to recording. Based on the depth of the probe tip, the angle of penetration, and the position of these sites, the location of units could be determined. Units recorded outside the target structure were excluded from analysis.

Extracellular voltages were amplified and digitized at 30 kHz via an RHD2164 amplifier board and acquired by an RHD2000 system (Intan Technologies). No filtering was performed at the data acquisition stage. Kilosort ^55^ was used for initial spike clustering. We configured Kilosort to highpass filter the input voltage time series at 300 Hz. The automatic clustering results were manually curated in Phy for putative single-unit isolation. We noticed a previously reported issue of Phy double counting a small fraction of spikes (with exact same timestamps) after manually merging certain clusters, thus duplicated spike times in a cluster were post-hoc fixed to keep only one.

Cluster quality was quantified using two metrics (Extended Data Fig. 3a-c,e). The first was the percentage of inter-spike intervals (ISI) violating the refractory period (RPV). We set 2.5 ms as the duration of the refractory period and used 1% as the RPV threshold above which clusters were regarded as multi-units. It has been argued that RPV does not represent an estimate of false alarm rate of contaminated spikes ^56, 57^ since units with low spike rates tend to have lower RPV while high spike rate units tend to show higher RPV even if they are contaminated with the same percentage of false positive spikes. Therefore, we estimated the contamination rate based on the reported method ^56^. A modification was that we computed a cluster’s mean spike rate from periods where the spike rate was greater than 0.5 spikes/s rather than from an entire recording session. As a result, the mean spike rate reflected more about neuronal excitability than task involvement. Any clusters with more than 15% contamination rate were regarded as multi-units. Combining these two criteria in fact classified less single-units than using a single, though more stringent, RPV of 0.5%. A low RPV can fail potentially well isolated fast spiking interneurons whose ISIs can frequently be shorter than the set threshold.

### Photostimulation

Bilateral stimulation of the brain was achieved using a pair of optic fibers (0.39 NA, 400 µm core diameter) that were manually positioned above the clear skull prior to the beginning of each behavioural session. These optic fibers were coupled to 470 nm LEDs (M470F3, Thorlabs). The illumination power was externally controlled via WaveSurfer (http://wavesurfer.janelia.org). Each stimulation had a 2 s long 40 Hz sinusoidal waveform with 0.1 s linearly modulated ramp-down at the end. The peak powers in the main experiments were 16 mW and 8 mW. We used the previously reported 50% transmission efficiency of the clear-skull preparation ^9^ and report the estimated average power in the Results. There was a 10% chance of light delivery triggered at each of the following points in a sequence: cue onset, the fourth (middle) touch, or the first touch after water delivery. To ensure that the light from photostimulation did not affect the mouse’s performance through vision, we set up a masking light with two blue LEDs directed at each of the mouse’s eyes. Each flash of the masking light was 2 s long separated by random intervals of 5-10 s. This masking light was introduced several training sessions in advance of photostimulation to ensure the light no longer affected the behaviour of the mouse. In addition, the optic fibers were positioned to shine light from ∼5-10 mm above the animal’s head on these days leading up to photostimulation.

In a subset of silicon probe recording sessions (related to Extended Data Fig. 2a-d), we used an optic fiber (0.3 NA, 400 um core diameter) to simultaneously photoinhibit the same (within 1 mm) or a different cortical region (∼1.5 or ∼3 mm away) via a craniotomy. The tip of the fiber was kept ∼1 mm away from the brain surface. For testing the efficiency of photoinhibition, the same 2 s photostimulation was applied but only at the mid-sequence, with 7.5% probability for each of the four powers (1, 2, 4 and 8 mW). For each isolated unit, the photo-evoked spike rate was normalized to that obtained during the equivalent 2 s time window without photostimulation. To avoid a floor effect, we also excluded units that on average fired less than one spike during the no stimulation windows. We classified units as putative pyramidal neurons if the width of the average spike waveform (defined as time from trough to peak) was greater than 0.5 ms, and as putative fast spiking interneurons if shorter than 0.4 ms or if units had more than twice the firing rate during 8 mW photostimulations than during periods of no stimulation.

### Histology

Mice were perfused transcardially with PBS followed by 4% PFA in 0.1 M PB. The tissue was fixed in 4% PFA at least overnight. The brain was then suspended in 4% agarose in PBS. A vibratome (HM 650V, Thermo Scientific) cut coronal sections of 100 μm that were mounted and subsequently imaged on a fluorescence microscope (BX41, Olympus). Images showing DiI and DiD fluorescence were collected in order to recover the location of silicon probe penetrations.

### Trial selection and alignment

The first trial and the last trial were always removed due to incomplete data acquisition. Trials in which mice did not finish the sequence before video recording stopped were excluded from the analyses that involved kinematic variables of tongue motion. The following procedures are specific to analyses related to spike PETH, linear regression and decoding.

Due to individual variability, different mice tended to lick at slightly different rates within lick bouts. The same mice might also perform a bit faster in one sequence direction than the other. Even in a given direction, one might start faster and then slow down a little, or slower first and faster later. When aligning trials from heterogeneous sources, a 10% difference in lick rate, for instance, will result in a complete mismatch (reversed phase) of lick cycle after only 5 licks. Therefore, prior to the analyses that are sensitive to inconsistent lick rates (Figs. 4, 5, 6 and Extended Data Figs. 3, 4, except for Fig. 4b,d,f), we linearly stretched or shrunk inter-lick intervals (ILIs) within each lick bout to a constant value of 0.154 s (i.e. 6.5 licks/s), which is around the overall mean. The timestamps of lick used to compute ILIs were contact onsets for touches or the time at *L_max_* for missed licks. A lick bout was operationally defined as a series of consecutive licks in which every ILI must be shorter than 1.5 × the median of all ILIs in the entire behavioral session. For ease of programming, we compensatorily scaled the time between the last lick of a trial and the start of the next trial to maintain an unchanged global trial time. Original time series, including spike rates and *L’*, were obtained prior to temporal alignment. After alignment, the behavioral and neural time series were resampled uniformly at 400 samples/s.

After temporal alignment, we used a custom clustering algorithm to find a group of trials with the most similar sequence performance. First, all trials of a given sequence in a behavioral session were collected and a time window of interest was determined. In Fig. 4 and Extended Data Fig. 3, we used the same time windows as the corresponding PETHs. In Fig. 5, we used −1 to 1 s from the 4th (middle) touch. In Fig. 6, we used −0.5 to 1 s from the first lick touching water. The duration of a time window is denoted as *T*. Second, for each trial, a feature vector was constructed which included 7×*T* lick onset times and 7×*T* touch onset times within the time window. Insufficient timestamps were padded with zeros. Finally, pairwise euclidean distances were computed among feature vectors of all candidate trials and we chose a subset of *N* trials with the lowest average pairwise distance, i.e. those that have the most similar lick and touch times. The number *N* was set to 1/3 of available candidate trials with a minimal limit of *N* = 10 trials. We used this relatively low fraction mainly to handle the greater behavioral variability in sequences with backtracking. To handle trial-to-trial variability in sequence initiation time (defined as the interval from the cue onset to first touch onset), which was not captured in our feature vectors, prior to clustering we limited trials to those with sequence initiation time less than 1 second.

### Behavioral quantifications

Learning curves included data from all trials except for the first and the last trial of each session. Values in the learning curves were averaged in bins of 100 trials, with 50% overlap of consecutive bins.

The duration of individual licks was variable. In order to average quantities within single licks (Figs. 1-3,6 and Extended Data Fig. 2), we first linearly interpolated each quantity using the same 30 time points spanning the lick duration (from the first to the last video frames of a tracked lick). *L’* was computed before interpolation. When the tongue was short, the regression network showed greater variability in determining *θ* and sometimes produced outliers. Thus, we only show *θ* when *L* is longer than 1 mm. In addition, any “lick” with a duration shorter than 10 ms was excluded.

The instantaneous lick rate was computed as the reciprocal of the inter-lick interval. The instantaneous sequence speed was defined as the reciprocal of the duration from the touch onset of a previous port position to the touch onset of the next.

The rate of licks, rate of touches, *Θ_shoot_* and *L_max_* as a function of time (Fig. 3c-f and Extended Data Fig. 2e-j,o,p) were computed using 0.2 s time bins. The summary quantifications (Fig. 3g-j and Extended Data Fig. 3k-n) used data averaged within 1 s after the start of photoinhibition (or the equivalent time in no-inhibition trials). The shorter window helped to minimize the effects “bleeding over” from mid-sequence to initiation, and from consumption to mid-sequence. Although this was not an issue for the consumption period, we nevertheless used the 1 s window for consistency. The summary metric SD(*Θ_shoot_*) was obtained by averaging the standard deviation of *Θ_shoot_* in each 0.2 s time bin within the 1 s window. Other metrics were directly computed without binning.

Directly averaging trials pooled across animals assumes that different animals follow the same distribution, and thus underestimates potentially meaningful animal-to-animal variability. To incorporate this variability, we performed a hierarchical bootstrap procedure ^58^ when computing confidence intervals where noted. In each iteration of this procedure, we first randomly sampled animals with replacement, then, from each of these resampled animals, sampled sessions with replacement, and then trials from each of the resampled sessions. The statistic of interest was then computed from each of these bootstrap replicates. In our optogenetic inhibition experiments, each animal only contributed one behavioral session for a given experimental condition. Therefore, the hierarchy only had two levels.

### PETH, NNMF and t-SNE

Spike rates were computed by temporal binning (bin size: 2.5 ms) of spike times followed by smoothing (15 ms SD Gaussian kernel). The smooth PETHs were computed by averaging spike rates across trials. Each unit has 6 PETHs: 3 time windows (for sequence initiation, mid-sequence and sequence termination) each in 2 standard sequences (left to right and right to left). We excluded inactive units whose maximal spike rate across the 6 PETHs was less than 10 spikes/s. For the rest, we normalized PETHs of each unit to this maximal spike rate. To construct inputs to NNMF and t-SNE, the 6 PETHs of each unit were downsampled from 2.5 ms per sample to 25 ms per sample and were concatenated along time to form a single feature vector.

NNMF was performed using the MATLAB function “nnmf” with default options. We empirically chose to compute 10 clusters as too few clusters tend to merge response patterns tuned to adjacent stages of sequences, whereas too many clusters provided little help in extracting the major response patterns from the data. NNMF is a close relative of principal component analysis (PCA) that has gained increasing popularity for processing neural data. The algorithm finds a small number of activity patterns (equivalent to principal components in PCA) along with a set of weights for each neuron, so that the original PETHs can be best reconstructed by weighted sums of those activity patterns. As a result, a small number of activity patterns (or dimensions) is usually able to capture the main structure of the original PETHs, and a neuron’s weights quantifies the degree to which its activity reflects each pattern. In the context of clustering, each pattern describes representative activity of a cluster, and the pattern with the greatest weight for a neuron determines its cluster membership (Extended Data Fig. 3k).

t-SNE is a nonlinear dimensionality reduction method. We used the MATLAB “tsne” function with customized options (’Algorithm’, ‘exact’, ‘Distance’, ‘cosine’, ‘NumDimensions’, 2, ‘Perplexity’, 50, ‘Standardize’, false) to find a 2-D embedding of the feature vectors (Extended Data Fig. 3g). t-SNE per se does not cluster data. Therefore, we fitted Gaussian mixture models with various numbers of components and used BIC to determine the optimal number of clusters (Extended Data Fig. 3h). Since the t-SNE results can be sensitive to initial conditions, we repeated the computation with 50 different seeds of a random number generator and obtained a median cluster number of 8 (Extended Data Fig. 3i). The results shown are from the first run with 8 clusters.

### Linear model and PCA

In each linear regression, the predictors were normalized spike rates of simultaneously recorded units and the response was one of the 5 behavioral variables (*L*, *L’*, *θ*, *D*/*TP* or *τ*). Predictors and responses were sampled at 400 Hz and were *not* averaged across trials. PCA was performed using the same normalized spike rates.

Both single- and multi-units were included. To obtain normalized spike rates, we divided the original spike rate by the maximum spike rate or 5 Hz, whichever was greater. We adopted this “soft” normalization technique ^43^ to prevent weakly firing units from contributing as much variance as actively firing units. Note that this normalization was only necessary for PCA and did not affect the goodness of fit, *R*^2^, of linear models.

*L*, *L’* and *θ* were directly available at 400 samples/s. However, these variables had values only when the tongue was outside of the mouth. Therefore, samples without observed values were either set to zero (for *L*) or excluded from regression (for *L’* and *θ*). *D* was defined as 1 if the sequence was from right to left and 2 if left to right. *τ* simply took sample timestamps as its values. *TP* (Extended Data Fig. 5) was the same as *D* but defined based on the upcoming sequence.

Predicting single responses with dozens of predictors is prone to overfitting. Therefore, we chose the elastic-net variant of linear regression (using MATLAB function “lasso” with ‘Alpha’ set to 0.1), which penalizes big coefficients for redundant or uninformative predictors. A parameter *λ* controls the strength of this penalty. To find the best *λ*, we configured the “lasso” function to compute 10-fold cross-validated mean squared error (cvMSE) of the fit for a series of *λ* values. The smallest cvMSE indicates the best generalization, i.e. the least overfit. We conservatively chose the largest *λ* value such that the cvMSE was within one standard error of the minimum cvMSE. For each model, we derived the *R*^2^ from this cvMSE and reported it in Figs. 5 and Extended Data Figs. 4,5.

Each of the 5 regressions resulted in a vector of coefficients comprising one coefficient for each unit, and a constant. Each coefficient vector and constant was used to predict or decode the corresponding behavioral variable from a vector (or, for multiple time points, a matrix) of population spike rates. We did not perform additional cross-validation in decoding because (1) 30% of the decoding for standard sequences (0.5 to 0.8 s in Fig. 5 and −1.3 to −1 s in Extended Data Fig. 5) was from new data; (2) all decoding in backtracking sequences and during consumption period was from new data; and (3) the model has been proven the best generalization via cross-validation when selecting *λ*. The term “coefficient vector” and “coding axis” are used interchangeably.

The percent variance explained (VE) by principal components was simply derived from the singular values. To compute VE by each coding axis, we first obtained its unit vector and projected population spike rates onto it. The variance of the projected values is Var(explained). The total variance, Var(total), of the population activity is the sum of variance of all units. Finally, VE equals Var(explained) / Var(total) × 100%.

### Canonical correlation

In each session, we computed the trial-averaged neural trajectories in the five dimensional coding subspace and trajectories in the PC subspace from standard sequences (−0.5 to 0.8 s from 4th touch onset; right to left and left to right trajectories were concatenated). Canonical correlations were computed using MATLAB “canoncorr” function between trajectory matrices with the same number of dimensions. *N* correlation coefficients (*r*) quantified the correlation between the activity in each pair of the *N* dimensions after transformation. The average *r* across the *N* values reflected the overall alignment between the two transformed trajectories.

---

**Extended Data Fig. 1 (related to Figs. 1 and 2).**
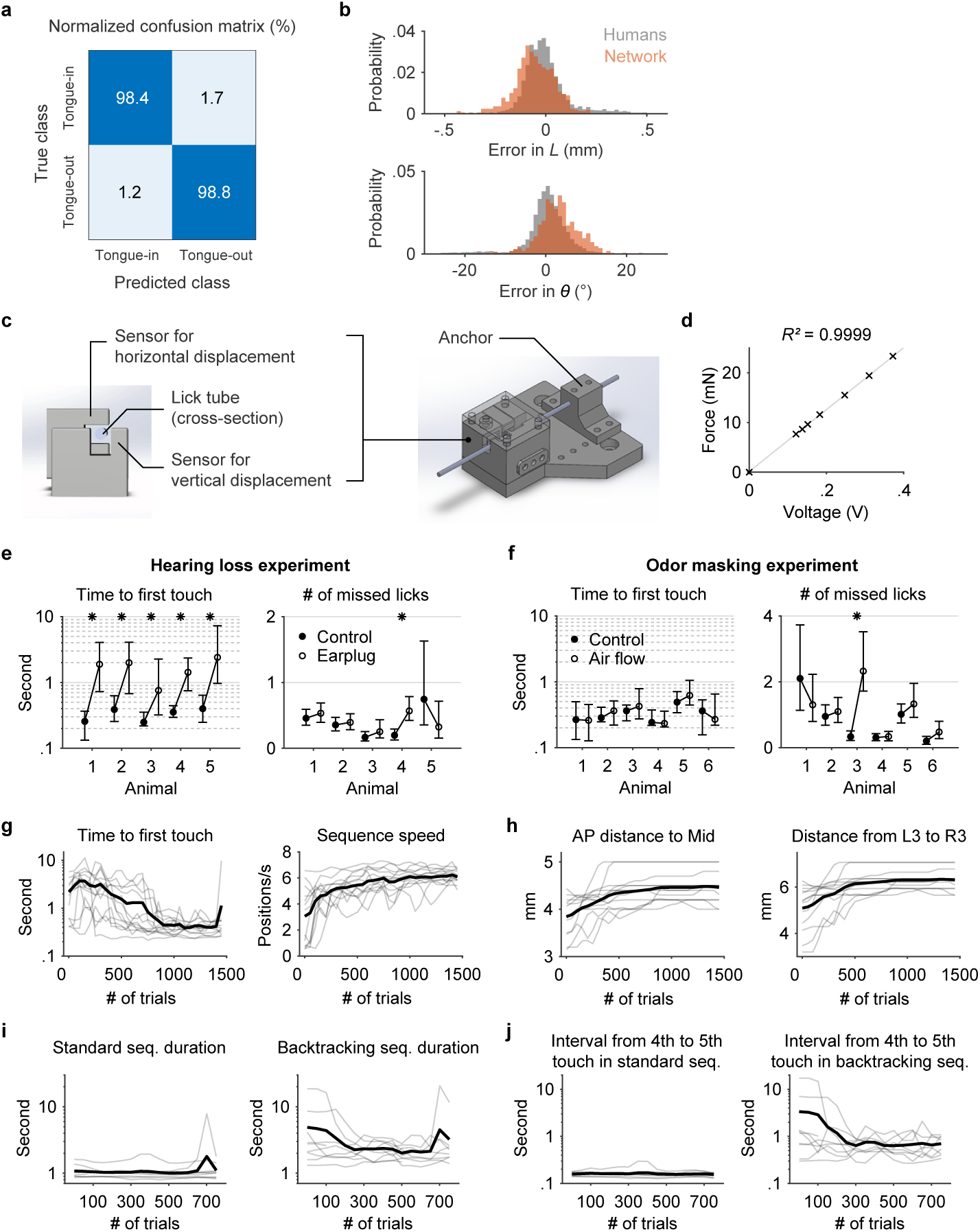
**a**, Confusion matrix showing the performance of the classification network. The numbers represent percentages within each (true) class (n = 1611 frames). **b**, Performance of the regression network. Top, the gray probability distribution shows how *L* from five human individuals varied from the mean *L* across the five. The red distribution shows how predicted *L* varied from the human mean. Bottom, similar quantification as the top but for *θ*. n = 573 frames. **c**, CAD images of the sensor core (left) and the assembly (right) with a lick tube. **d**, Linear relationship between the applied force and the sensor output voltage. **e**, Time to first touch (left; median ± interquartile range) and number of missed licks during sequence performance (right; mean ± 95% bootstrap confidence interval) in control versus hearing loss (earplug) conditions. ∗ p < 0.01, not significant otherwise, one-tailed KS-test, Bonferroni correction for multiple comparison. **f**, Time to first touch (left; median ± interquartile range) and number of missed licks during sequence performance (right; mean ± 95% bootstrap confidence interval) in control versus odor masking (air flow) conditions. ∗ p < 0.05, not significant otherwise, one-tailed KS-test, Bonferroni correction for multiple comparison. **g**, Learning curves for 13 individual mice (gray) and the mean (black) showing the reduced sequence initiation time (left) in response to the auditory cue and the increased sequence speed (right). **h**, Gradual increase in task difficulty (Methods) accompanying the improved performance shown in (**g**). **i**, Learning curves for 9 individual mice (gray) and the mean (black) showing the duration of time spent to perform standard (left) and backtracking (right) sequences. **j**, Similar to (**i**) but limited to the interval following the 4th (middle) lick in standard (left) or backtracking (right) sequences.

**Extended Data Fig. 2 (related to Fig. 3).**
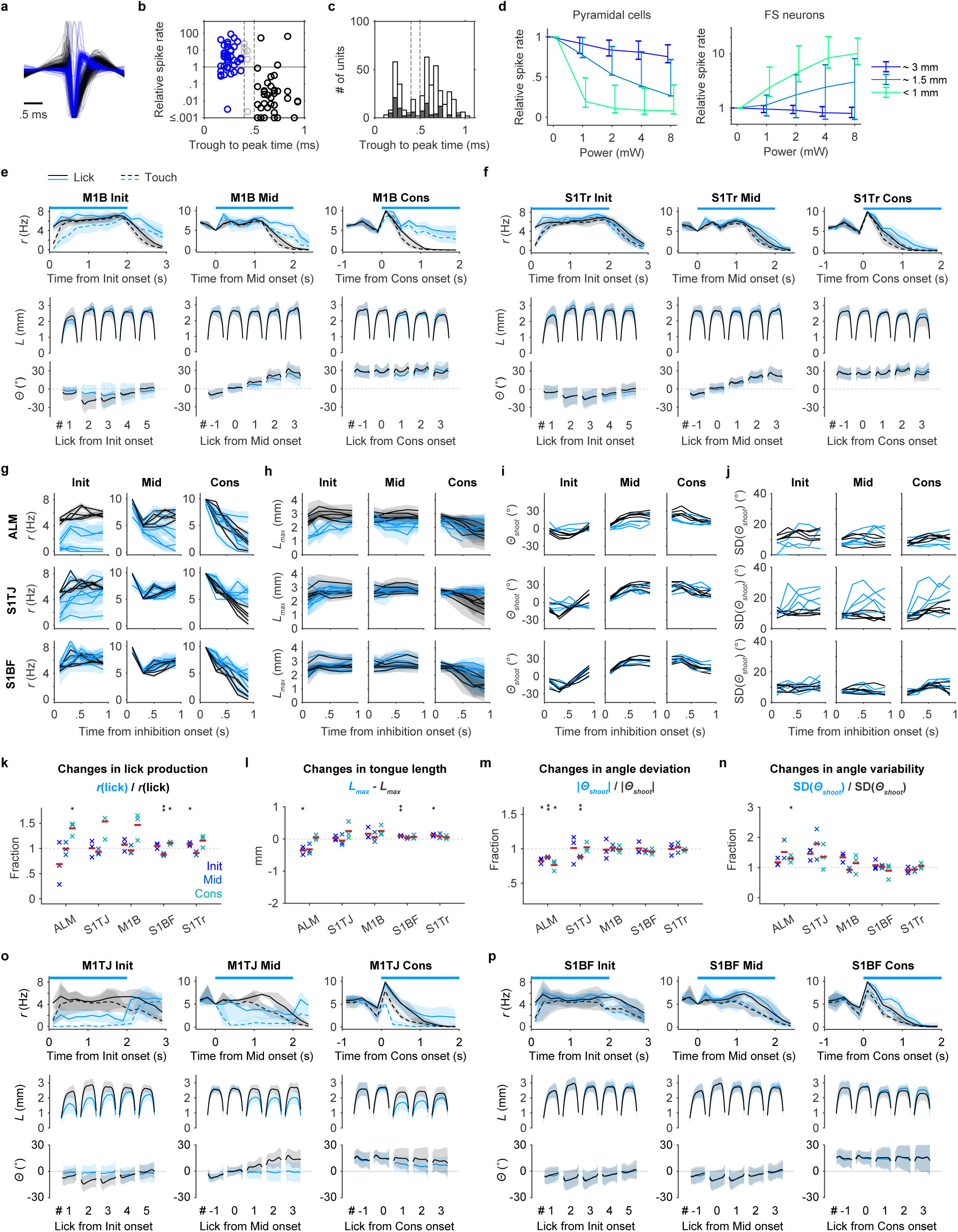
**a**, Average spike waveform of putative pyramidal cells (black; n = 224) and putative FS neurons (blue; n = 117), normalized to the amplitude of negative peaks. **b**, Relationship between spike widths (defined as the trough to peak time of average waveform) and changes in mean spike rate under opto illumination (4mW, within 1 mm) relative to baseline. Pyramidal cells (black; n = 42) and FS neurons (blue; n = 41) were classified by the two thresholds (dashed lines at 0.4 and 0.5 ms) with ambiguous units (gray; n = 6) in the middle. **c**, Distributions of spike widths from neurons in (**b**) (filled bars; n = 89) and from all neurons (empty bars; n = 414) including those where illuminations were not at recording sites. Classification thresholds are shown in dashed lines. **d**, Left, inhibition efficiency of putative pyramidal cells as a function of light power and distance away from the center of illumination (n = 224 units total). Right, similar to left but shows the excitation efficiency of putative FS neurons (n = 117 units total). Mean ± 95% hierarchical bootstrap confidence interval. **e,f**, Same as Fig. 3c but for M1B (n = 1240 trials) and S1Tr (n = 1269 trials), respectively. **g**, Rate of licks (*r*) of individual mice (mean ± 95% bootstrap confidence interval) with (blue) or without (black) inhibition in ALM (top), S1TJ (middle) or S1BF (bottom) and in each of the three periods. **h**, Similar to (**g**) but quantifies *L_max_* as a function of time for individual mice (mean ± SD). **i**, Similar to (**h**) but quantifies effects in *Θ_shoot_*. The standard deviation of *Θ_shoot_* (SD(*Θ_shoot_*)) is separately plotted in (**j**) for clarity. **k-n**, Same as Fig. 3g-j but with half of the inhibition power (2 mW). **o**, Same as (**e**) but for M1TJ. n = 709 trials from 3 mice. **p**, Same as (**e**) but for S1BF. n = 766 trials from the same 3 mice in (**o**).

**Extended Data Fig. 3 (related to Fig. 4).**
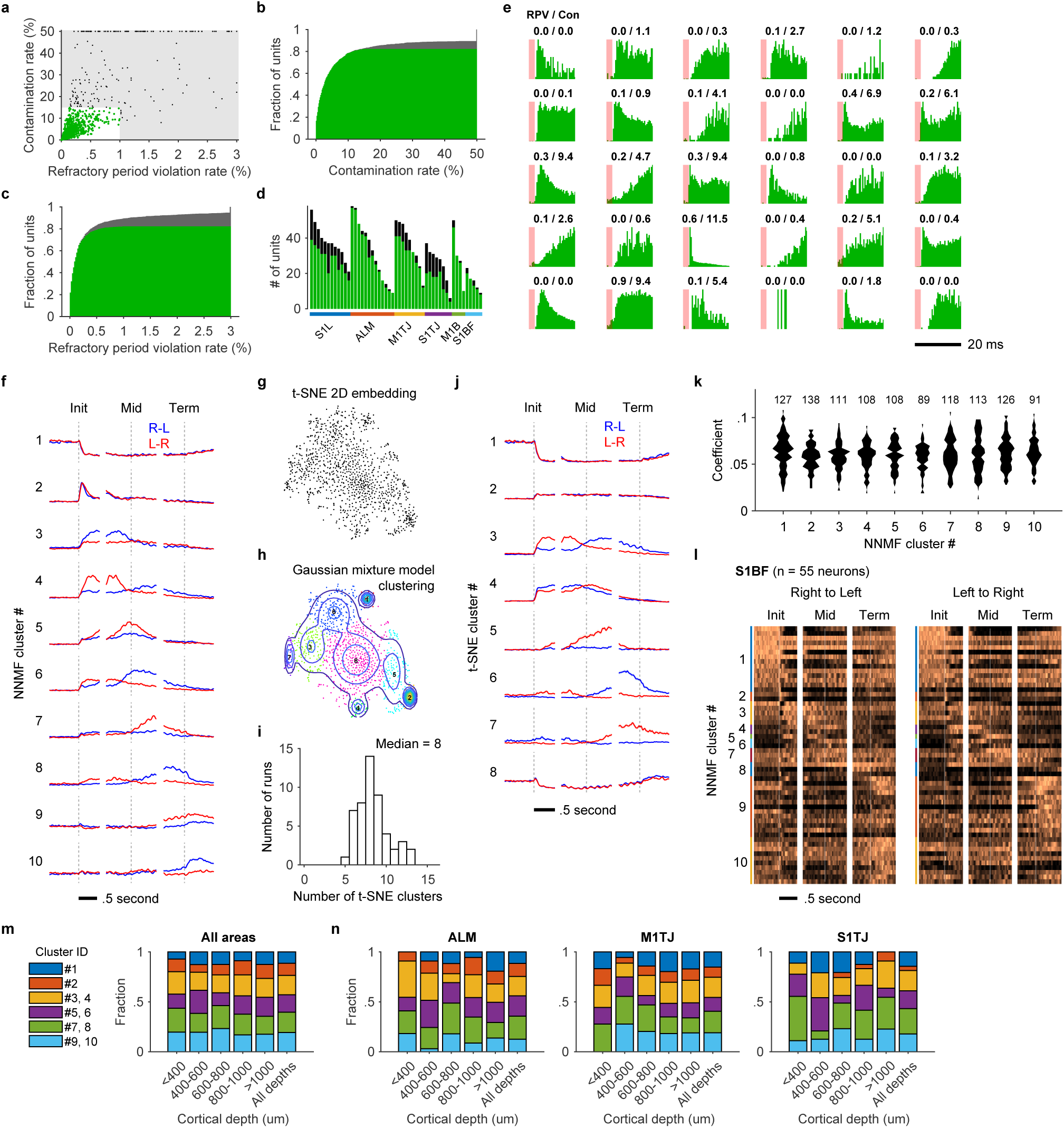
**a**, Contamination rates and refractory period violation rates of all recorded single-(green) and multi-units (black). The shaded region shows the thresholds for assignment as multi-vs single-unit. **b**, CDF of contamination rate including single-(green) and multi-units (gray). **c**, Same as (**b**) but for refractory period violation rate. **d**, The number of single-(green) and multi-units (black) recorded in each session, grouped by brain area. **e**, ISI histograms of randomly selected single-units. Refractory period violation rates (RPV) and contamination rates (Con) are labeled on the top (in percents). **f**, NNMF components that represent each of the ten PETH clusters. Right to left (blue) and left to right (red) activities (mean ± 95% bootstrap confidence interval) are overlaid together. The vertical lines are located at time zero in each period. The height of the lines represents the scale of normalized neuronal activity from 0 to 1. **g**, Two-dimensional embedding of neuronal PETHs via t-SNE. **h**, Nine clusters were fitted using a Gaussian mixture model. **i**, Distribution of the number of clusters computed with 50 different random number seeds. **j**, Same as (**f**) but showing the nine clusters computed via t-SNE and Gaussian mixture model. **k**, Distribution of membership coefficients for each of the NNMF clusters. The number of neurons in each cluster is labeled at the top. **l**, Same as Fig. 4c,e,g but for neurons in S1BF. **m**, Proportions of neurons (n = 1312 in total) from different clusters at different cortical depths. Some clusters were grouped together (e.g. 3 and 4) since they fired around the same time but only differed in direction selectivity. **n**, Similar to (**m**) but broken down for ALM (n = 334), M1TJ (n = 237) and S1TJ (n = 118).

**Extended Data Fig. 4 (related to Figs. 5).**
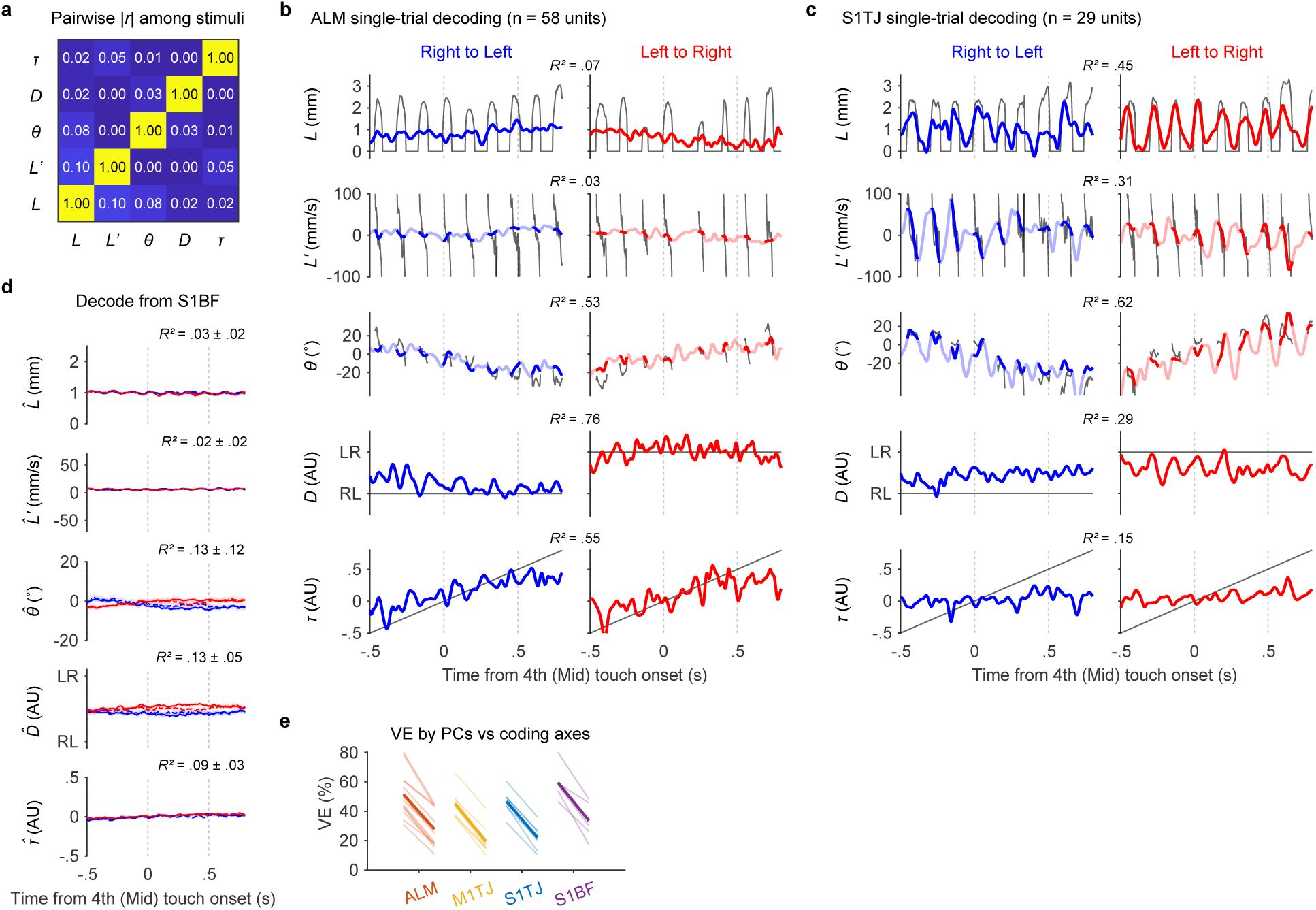
**a**, Absolute mean pairwise Pearson’s correlation coefficients among the five behavioral variables. (n = 35 sessions) **b**, Single-trial decoding of the five behavioral variables (rows; black traces) from 58 simultaneously recorded ALM units in a right to left (left) and a left to right (right) sequence. **c**, Same as (b) but decoding from 29 simultaneously recorded units in S1TJ. **d**, Same as Fig. 5b but for decoding from S1BF (n = 5 recordings). **e**, Total percent variance explained (VE) by the first five principal components (left in each region) versus that by the five coding axes (right in each region) during sequence execution. Lighter lines show individual recording sessions and thicker lines show the means.

**Extended Data Fig. 5.**
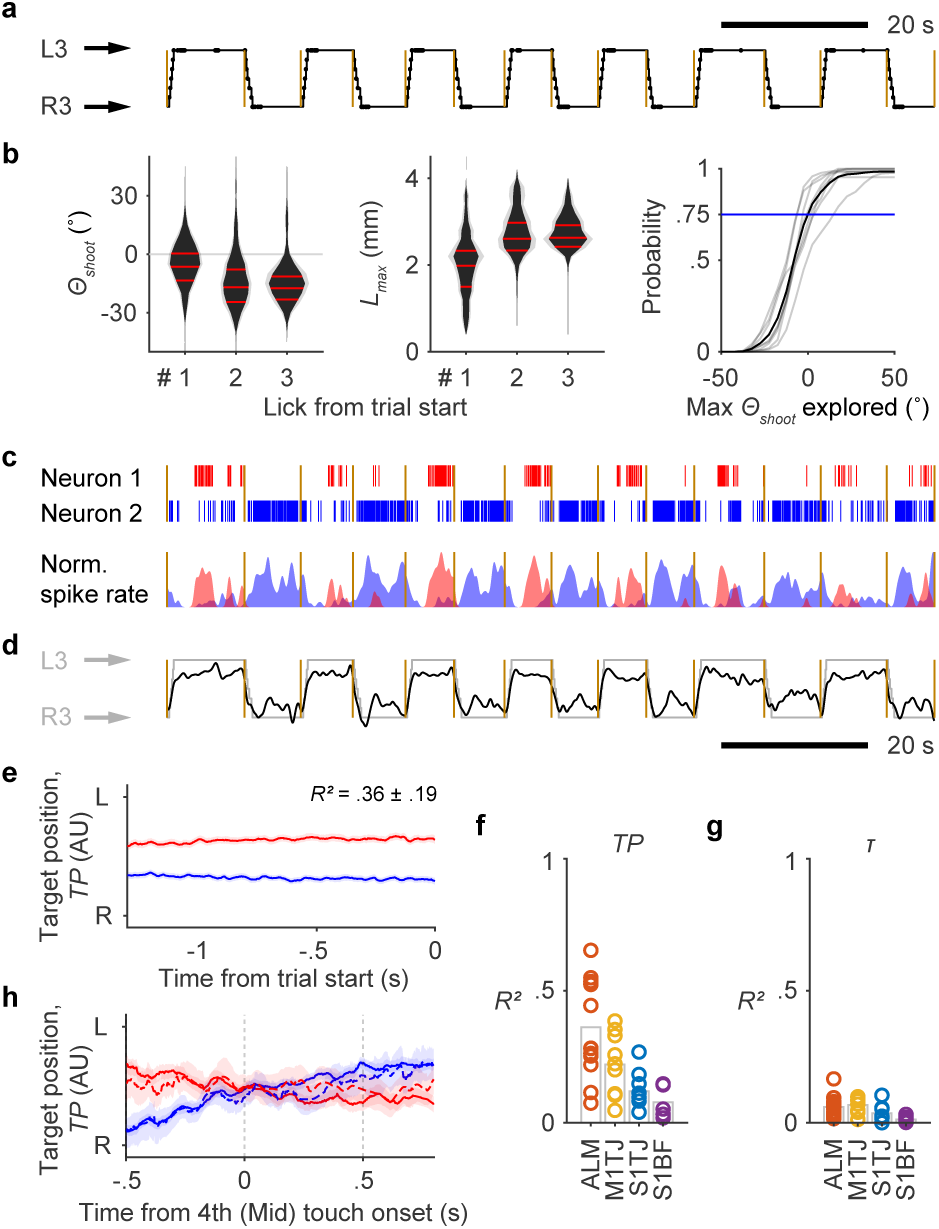
Coding of Upcoming Sequences in ALM. **a**, Depiction of sequences performed by a mouse in alternating directions across 14 consecutive trials. Trial onsets are marked by yellow lines. Port positions shown in the black trace are overlaid with touch onsets (dots). **b**, Probability distributions of *Θ_shoot_* (left) and *L_max_* (right) for the first 3 licks at the start of a sequence (n = 8 mice; mean ± SD). The negative y-axis points to the side at which the port is located. The CDF (8 individual mice in gray and the mean in black) of the maximal *Θ_shoot_* explored before touching the port (at the side of negative *Θ*). The blue line shows the probability of successfully locating the port without exploring beyond the midline. **c**, Top, rasters of two example units which had persistent and target position (*TP*) selective firing during the 14 consecutive trials in (**a**). Bottom, normalized and smoothed (0.25 s SD Gaussian kernel) spike rates of the two units. **d**, Decoded instantaneous *TP* (dark trace) from 58 simultaneously recorded units in ALM, overlaid with normalized port position (light trace). **e**, Decoding of *TP* from ALM (mean ± 99% bootstrap confidence interval) before upcoming right to left trials (blue) or left to right trials (red). Cross-validated *R*^2^ is shown (mean ± SD; n = 13 sessions). **f**, Goodness of fit for linear models that predict *TP* during ITIs, quantified by cross-validated *R^2^*. **g**, Same as (**f**) but for *τ*. **h**, Using the same linear models in (**d**) to decode *TP* during sequence execution. The decoded *TP* was averaged across trials (mean ± 99% bootstrap confidence interval) with standard sequences (solid lines) or backtracking sequences (dashed lines), each of which is either right to left (blue) or left to right (red).

Extended Data Table 1

Extended Data Table 1
| Animal ID | Genotype | Date of birth | Sex | Start age (mo) | Experiment length (mo) | Training history |
| --- | --- | --- | --- | --- | --- | --- |
| MX170903 | WT | 11/13/17 | M | 3 | 1 | SS |
| MX180201 | WT | 12/18/17 | M | 3 | 1 | SS |
| MX180202 | WT | 12/18/17 | M | 4 | 1 | SS |
| MX180203 | WT | 12/18/17 | M | 5 | 1 | SS |
| MX180601 | WT | 12/18/17 | M | 6 | 1 | SS, SS-SB (4), SS-BS-SB (3) |
| MX180602 | WT | 12/18/17 | M | 6 | 1 | SS |
| MX180701 | WT | 12/18/17 | F | 8 | 1 | SS, SS-BS-SB (4), SS-SB (2), SS-BS-SB (5) |
| MX180803 | VGAT-Cre+/-;Ai32+/- | 12/7/17 | M | 7 | 5 | SS, P'P' (1), R'R' |
| MX180804 | VGAT-Cre+/-;Ai32+/- | 12/7/17 | M | 7 | 5 | SS, P'P' (1), SS-BS (7), R'R' |
| MX181002 | VGAT-Cre+/-;Ai32+/- | 5/15/18 | F | 4 | 4 | SS, P''P'' (1), R'R' (5), SS (2), RR |
| MX181003 | VGAT-Cre+/-;Ai32+/- | 5/15/18 | F | 4 | 7 | SS |
| MX181101 | VGAT-ChR2-EYFP+/- | 7/29/18 | M | 5 | 3 | SS, PP (1), SS (2), PP (1), SR (1), RS (1), RR (3) |
| MX181301 | VGAT-ChR2-EYFP+/- | 7/29/18 | F | 5 | 5 | SS, PP (1), SS (2), PP (1), RS (1), RR (1), SS (3), RR |
| MX181302 | WT (VGAT-ChR2-EYFP-/-) | 7/29/18 | F | 5 | 1 | SS, PP (1), SS (2), PP (1), RS (3), RR |
| MX190101 | VGAT-Cre+/-;Ai32+/- | 7/9/18 | M | 9 | 1 | SS, PP (1), RR |
| MX190103 | VGAT-Cre+/-;Ai32+/- | 7/9/18 | M | 9 | 2 | SS, PP (1), RR |
| MX190201 | VGAT-Cre+/-;Ai32+/- | 12/30/18 | M | 5 | 4 | SS, RR |
| MX190401 | VGAT-Cre+/-;Ai32+/- | 2/22/19 | M | 6 | 1 | SS |
| MX190402 | VGAT-Cre+/-;Ai32+/- | 2/22/19 | M | 6 | 1 | SS |
| VC010102 | TH-Cre 1 | 4/8/19 | M | 6 | 2 | SS, RR |
| VC010103 | TH-Cre 1 | 5/5/19 | M | 5 | 1 | SS, RR |
| WO010401 | Avil-Cre | 6/18/19 | F | 2 | 3 | SS, RR |
| WO010402 | Avil-Cre | 6/18/19 | M | 2 | 3 | SS, RR |
| MX190602 | VGAT-Cre+/-;Ai32+/- | 6/3/19 | M | 5 | 1 | SS, RR |

